# A food chain of ribozymes built on an ancestral strand

**DOI:** 10.64898/2025.12.22.695801

**Authors:** R Weiss, C Farage, N Mellul, A Levy-Zamir, N Joseph, Y Svirskaya, O Ben Dov, N Vernia, S Ittah, A Ionescu, D Ionescu, I Bachelet

## Abstract

Molecular life forms in the prebiotic earth were capable of self-reproduction from basic building blocks. However, scarcity of building blocks likely led to the evolution of heterotrophic, predatory species. In this work we reconstruct a simple model of such interactions between RNA molecules, based on RNA strands found in sinkhole mineral pools in the Dead Sea. These RNA strands exhibited significant stability in this divergent chemical environment, and contained sequence motifs reminiscent of two ribozyme species, suggesting they could be ancestral to both. We built a system in which a single ancestral RNA strand serves as a limiting resource forming two different species of ribozymes that compete over it. One of these species is capable of metabolizing the other in order to free more of this resource, driving the assembly of additional copies of itself. This interaction resembles a rudimentary food chain, the dynamics of which is sensitive to environmental factors such as temperature and magnesium concentrations, with competitive advantage shifting between the species. Taken together, these findings highlight a hypothetical feature of prebiotic life, and hint at the possibility that heterotrophic RNA is a phenomenon that might still occur today in some unique geochemical niches, and even in the overlooked background of modern biological systems.

## Introduction

Prebiotic worlds are likely inhabited by molecules exhibiting life-like features, such as sensing and responding to environmental cues, metabolic activity, self-organization, and self-reproduction. Moreover, these proto-organisms likely interact with each other in ways similar to those constructing modern ecological systems.

The RNA world hypothesis ^1–3^ is generally accepted as one of the leading theories on the origin of life. Unlike DNA and proteins, RNA functions in nature both as a genetic message and chemical catalyst. These two functions pose contradicting sets of features, which the chemistry of RNA appears to satisfy almost to an optimum^2^.

The improbability of prebiotic formation of RNA nucleobases and oligomers in the primitive earth has been previously discussed in a geochemical context ^4–8^. Alternative RNA nucleobases have been proposed as putative precursors to RNA, and their behavior and stability have been studied ^9,10^. Regardless of the exact original mechanism that brought RNA into existence, the RNA world hypothesis implies that, at some later point in time, RNA molecules existed at sufficient concentrations enabling them to collide, interact, and recombine. These interactions mark a path for the emergence of structural and functional complexity ^11,12^, which is arguably more plausible than the emergence of complex species composed of a single long RNA molecule. The latter has been the aim of several previous experiments aiming to produce self-polymerizing RNA enzymes (ribozymes) by *in-vitro* evolution, and although some of these studies produced single-molecule ribozymes with impressive catalytic performance ^13,14^, it is generally assumed that ancient RNA-polymerizing ribozymes had low processivity, and were therefore likely capable of synthesizing only relatively short RNA oligonucleotides^12^.

The proto-organisms of the RNA world were likely capable of self-reproducing from basic building blocks, although the exact nature of these blocks is unknown. Intriguingly, while modern RNA polymerases use single RNA nucleobases as building blocks, the primitive blocks might in fact have been short RNA oligomers. Some naturally occurring mineral catalysts drive synthesis of RNA oligomers ^6,15^, and several ribozymes were produced *in-vitro* using RNA parts as building blocks ^16–19^, highlighting this possibility. RNA oligomers as short as 5 bases are capable of catalytic activity ^20,21^, blurring the distinction between basic building blocks and functional ribozymes in the RNA world.

The small RNA parts that interacted and self-organized to form the first ancient life-like RNA complexes were hypothetically poured into the environment ^21,22^. However, this constant flux of RNA fragments, which produced the first RNA life forms, might have become insufficient at later stages, as stable forms emerged that utilized these parts with increasing efficiency. An outcome of this hypothetical scenario is competition between different RNA species for the limiting parts required for their reproduction. Such competition may have driven the emergence of the first predatory species of RNA molecules ^23^, which could break down other RNA species in order to free parts for their own reproduction. Our aim in this work was to build on this hypothesis and explore possible designs of predatory RNA interactions.

Several observations on ribozymes are interesting in the context of our hypothesis. First, naturally occurring ribozymes ^24^ are capable of diverse features, some of which are directly life-like, such as the ability to respond to metabolite concentrations ^25^. Second, the basic metabolic operations, cleavage and assembly, are the main functions exhibited by known natural ribozymes that operate on RNA, which are either nucleases, or ligases, or both ^26^. RNA-polymerizing ribozymes are known to date from *in-vitro* evolution experiments. Third, although in their biological contexts, modern natural ribozymes are typically self-cleaving, they are capable of cleaving in *trans*, demonstrating that this is a valid mode of ribozyme action. It is conceivable that the self-cleaving ribozymes existing today as integral parts in genomes and transcripts, are the “fossilized” remains of these once-free RNA organisms. Fourth, functional ribozymes can be recombined from short RNA molecules ^11,27^. Taken together, these observations support the hypothesis that ancient ribozymes could operate on other ribozymes, break them down to simpler parts, and assemble them into functional ribozymes, forming molecular equivalents of food chains.

We were interested in reconstructing a plausible proof of existence for such interactions, using simple ribozymes as a starting point. The hammerhead ribozyme, for example, has several attractive properties in this regard. First, it is found across all kingdoms of life from viroids ^28^ to humans ^29^, and in multiple topological variations ^30–32^, suggesting that this structure has ancient origins. Second, it can cleave, ligate ^33^, and operate in *trans* ^34,35^. Third, it is a relatively small structure, thus arguably has a higher likelihood of formation compared with large, complex ribozymes. Fourth, although the modern hammerhead ribozyme is a single strand, it is also active as a heterodimer of fragments ^36^, and can be assembled successfully from short parts ^27^. The hairpin ribozymes share most of these properties as well ^37^ ^11,27^. The hammerhead and hairpin ribozymes have been previously shown to operate on each other artificially ^27^.

We have recently conducted a survey of environmental RNA from mineral pools in the Dead Sea, and isolated a group of RNA strands that contain sequence motifs common to both hammerhead and hairpin ribozymes. Thus, these molecules are capable of serving as scaffolds for building both ribozymes, suggesting they may have been ancestors of both. These sequences were interesting in other aspects as well, for example being composed of *k-*mers that showed significant stability in Dead Sea water.

In present work, we utilized these RNA molecules to build a system in which a hammerhead and hairpin ribozymes play two competing, RNA-based, heterodimeric “organisms”. They were assigned the roles of predator and prey, respectively, with the prey renewing by environmental flux. We demonstrate that the competition leads effectively to reproduction of the predatory species and identify conditions that give rise to alternative dynamics and outcomes. We propose that such food chains as a plausible phenomenon in the RNA world. Moreover, these observations highlight the possibility that, in addition to being a Darwinian force in the RNA world, heterotrophic RNA is an extant phenomenon, and might occur in some unique, isolated, and chemically divergent geochemical niches.

## Results

The Dead Sea **(Fig. 1A, 1B)** is a hypersaline terminal desert lake located at the borders of Jordan, the Palestinian Authority and Israel. The total dissolved salts concentration in the lake is approximately 350 g L^-1^, with high concentration of the divalent cations Mg^2+^ and Ca^2+^ (2 M and 0.5 M, respectively, **Fig. 1C**), which play a major role in making the Dead Sea more hostile than other hypersaline environments. The Dead Sea has been isolated from the Mediterranean Sea for over 2 million years, reaching its overall current shape ca. 11,000 years BCE ^38^. Currently, the lake consists of a deeper northern basin (deepest point at ∼725 m below sea level) and a southern basin, which has dried out in the last decades and is artificially filled in for commercial purposes. The Dead Sea was a meromictic lake with hypersaline, anoxic and sulfidic deep waters and a seasonally varying mixolimnion until 1979 ^39^. However, the decreasing input of water into the lake since the beginning of the 20th century led to a continuous decrease in the water level ^39,40^ resulting in increased salinity and the eventual overturn of the lake in 1979. Life in the Dead Sea has been documented since the 1930s ^41^. Blooms of the halophilic green alga *Dunaliella* followed by high densities of heterotrophic Archaea have been documented three times between 1964 and 1992 ^42^ and never since. Recently, highly localized dense microbial communities were documented in the vicinity of freshwater underwater springs in several locations on the Dead Sea shore ^43,44^.

**Figure 1.**
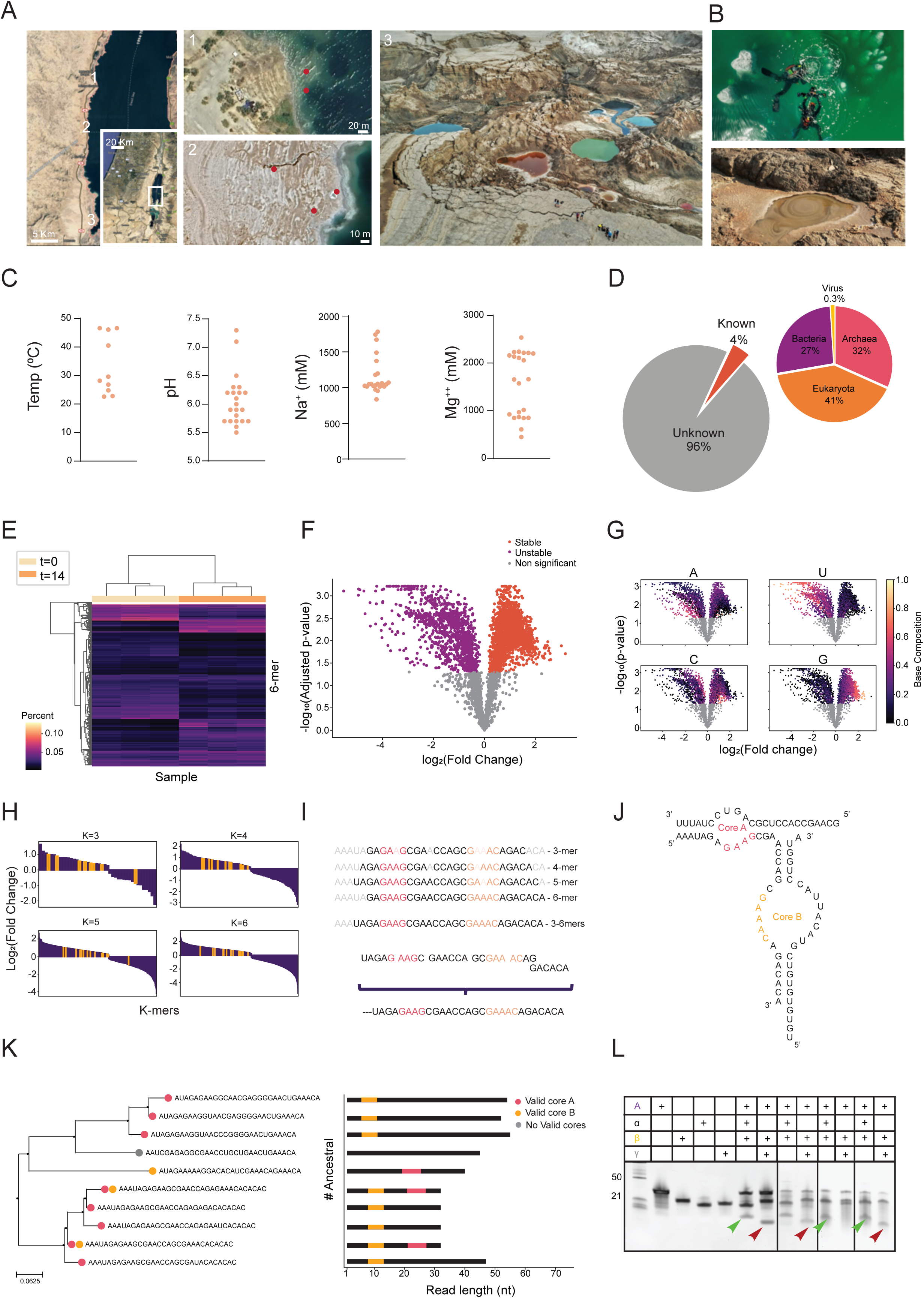
Sampling in the dead sea and Analysis of collected samples. **A,** A detailed map highlighting the specific sampling locations within the Dead Sea region. Red dots indicate dive sites (label 1) where samples were collected, sites of hot spring samples (label 2), and an illustrative photo of the pools sampled (label 3), providing a geographic overview of the research scope. **B,** Imagery related to the sampling process: an upper photo shows divers in the Dead Sea engaged in sample collection, while a lower photo captures one of the pools from which samples were taken, offering a visual insight into the fieldwork conditions. **C,** Displays a series of graphs detailing the temperature (Temp), pH, sodium (Na++), and magnesium (Mg2+2+) levels measured across various samples. **D,** the left panel reveals that the majority of sequences derived from Dead Sea collections were novel, indicating significant unexplored biodiversity. The right panel further classifies the known sequences according to their biological kingdom. **E,** Hierarchical clustering heatmap for K-mers percentage demonstrating clustering of samples according to condition. Bright and dark colors show higher and lower percent of K-mer in the sample, respectively. Yellow and orange labels indicate time points**. F,** Volcano plot of K-mer stability in Dead Sea sample at t=0 and t=14 days of exposure to Dead Sea water. Depicts Log2 Fold Change on the x-axis and adjusted p-value for multiple testing (FDR) on the y-axis. Each K-mer is represented as a dot, red dots are significant stable k-mers, purple dots are unstable k-mers. **G,** K-mer stability volcano plot colored according to base composition. Brighter color indicates higher proportion of base A, U, C or G within the K-mer.**H,** Bar plots of significantly different K-mers (K=3,4,5,6), sorted by descending order of Log2 Fold Change. Highlighted in yellow are k-mers corresponding to the ancestral strand. **I,** Ancestral strand assembly from significant stable K-mers. On top is the ancestral strand sequence coverage for K=3,4,5,6 from significantly stable K-mers from Dead sea water. Below is a hypothetical assembly route of the Engine sequence. Highlighted in orange are the two catalytic cores essential for the ribozymes activity. The K-mers cover 90.625% of the engine’s sequence, which is higher than the coverage of a scrambled (p-value= 0.0094) or random (p-value= 0.00028) engines, according to one-sample t-test, n=30. **J,** a schematic representation of the HH ribozyme’s design, with particular emphasis on the critical regions: core A and core B. **K,** Ancestral strands in Dead Sea. On the left, a phylogenetic tree constructed from BLAST hits of the ancestral strand identified in Dead Sea samples. On the right are the hits highlighted within their position in the read they derived from. Colors indicate core integrity. **L** Gel electrophoresis results that compare the cleavage activity of found ribozymes under laboratory conditions (left rectangle) against those observed in three distinct Dead Sea samples (small rectangles). The green arrow highlights cleavage products indicative of HH activity, while the red arrow points to those resulting from HP activity, showcasing the experimental validation of ribozyme function in varied environmental conditions.

Free RNA was isolated from various locations within the Dead Sea and the surrounding sinkhole lakes, and sequenced. Approximately 4% of the RNA was mapped to known organisms, particularly halophilic and thermophilic sulfate-reducing organisms **(Fig. 1D, Fig. S1)**. In order to identify specific enrichment patterns due to stability, we first measured the stability distribution of RNA sequences in Dead Sea water. To this end, an 18 nt random RNA library (1 µmol, equivalent to 4^18^ [∼ 10^11^] specific sequences ✕ ∼10^5^ copies per sequence) was incubated in independent triplicates in pure Dead Sea water for 14 days and sequenced **(Fig. 1E)**, and analyzed the most stable sequences which were enriched **(Fig. 1F)**. We found a significant trend of base frequency in the most stable sequences as follows: G>C>A>U **(Fig. 1G)**. We were specifically interested in identifying ribozyme cores in the data, and indeed interestingly within these samples, we detected a group of RNA molecules with sequence motifs that are common to both hammerhead and hairpin ribozymes, namely, the GAAA part of both hammerhead and hairpin cores, and the GAAG part of the hairpin core. These sequences, specifically 3, 4, 5, and 6-mers of them, were among the most stable fraction identified **(Fig. 1H, 1I)**. The data contained several variants of these strands **(Fig. 1J, 1K)**, which we termed “ancestral” due to their potential to give rise to both ribozymes. These sequences were shown experimentally to form catalytically-active ribozymes under standard reaction (laboratory) conditions and in the specific Dead Sea environments in which they were identified **(Fig. 1L, Fig. S2)**. These results inspired a design of putative “food chain” of ribozymes, where the ancestral strand is a limiting source for building two competing ribozyme species.

To create a system where two ribozyme species compete over the same limiting resource, enabling one ribozyme to break down the other one, a hammerhead and hairpin ribozymes were designed as follows. First, each ribozyme was fragmented into 2 separate RNA strands, one strand is the ancestral, and the second is a scaffold strand. Second, the ancestral strand sequence was redesigned such that the same strand could function in both ribozyme types, maintaining the residues that are critical for their function. Third, the hammerhead ribozyme was designed such that the hairpin scaffold strand could be used as the hammerhead substrate. The hairpin ribozyme was designed to operate on an unrelated substrate strand. Fourth, fluorophore-quencher pairs were used to tag the ribozyme parts to provide for a real-time reporter system for assembly of the ribozyme complexes **(Fig. 2A, 2B)**. Both ribozymes were tested separately to ensure they function properly and to measure their activity **(Fig. 2C, 2D)**.

**Figure 2.**
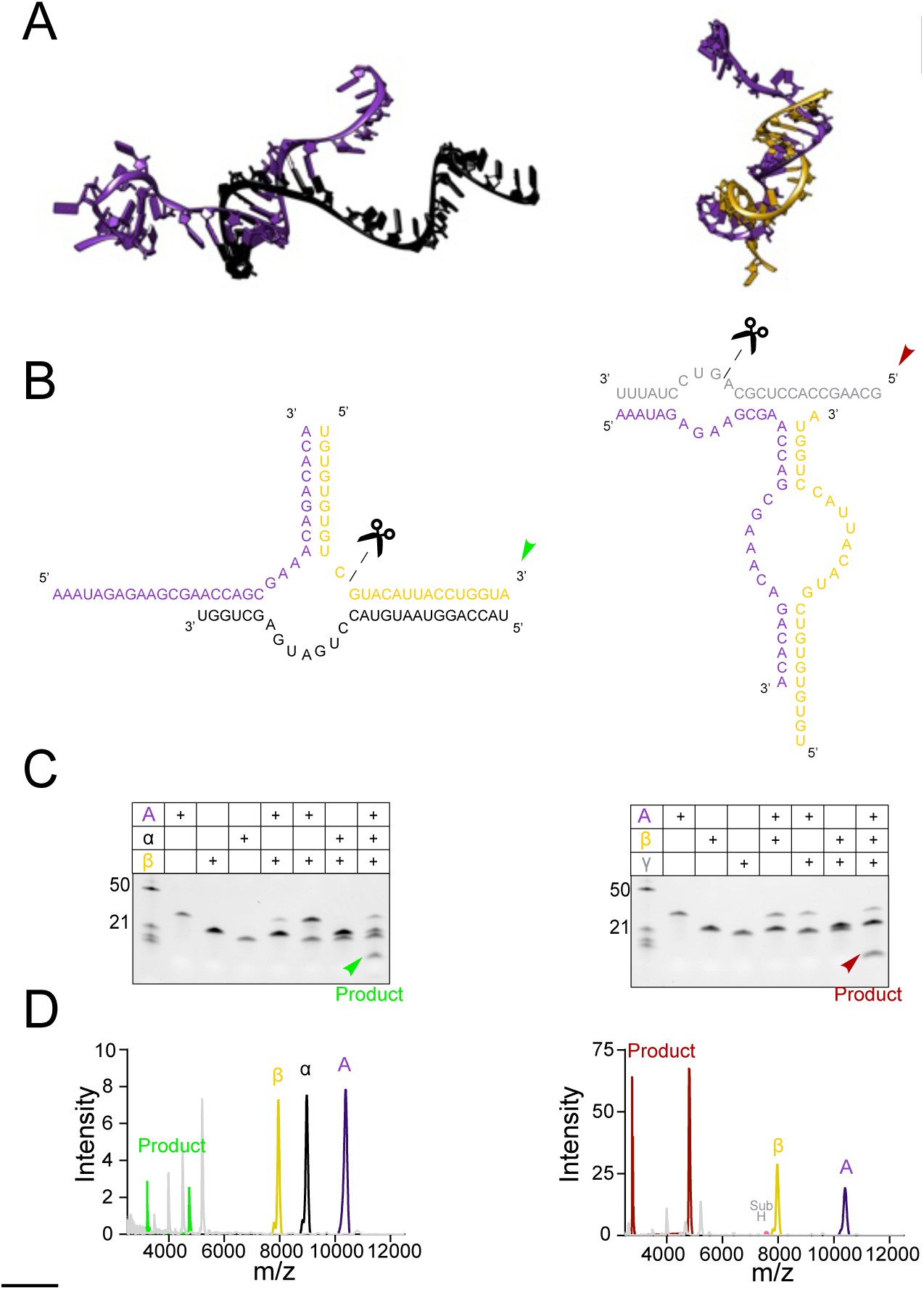
Food chain design and ribozyme function. **A,** 3D structural predictions of the HH and HP ribozymes. The HH ribozyme, depicted on the left, is formed by the interaction between the ancestral strand (colored in purple) and the α strand (black). The HP model, shown on the right, consists of the ancestral strand (purple) and the β strand (gold), illustrating the distinct structural components contributing to each ribozyme’s architecture. **B,** Detailed schematic of the design and substrate interaction for both ribozymes. The HH ribozyme (left schematic) incorporates three key elements: the ancestral strand (purple), the α strand (black), and its specific substrate, the β strand (gold). Conversely, the HP ribozyme (right schematic) is composed of the ancestral strand (purple), the β strand (gold), and its corresponding substrate, the γ strand. Scissors symbols indicate the cleavage sites on the substrates. **C,** Gel electrophoresis demonstrating the cleavage activity of the HH (left gel) and HP (right gel), arrows mark the cleavage product. **D,** Maldi-Tof analysis graphs for both HH (left graph) and HP (right graph) ribozymes. These analyses confirm the molecular outcomes of the cleavage reactions, providing precise mass spectrometric evidence of the expected cleavage products.

Next, the components were mixed in a single reaction, with excess of both scaffold strands over the engine strand as described in the methods section, and the reaction was allowed to proceed. We observed an increase in the number of hammerhead ribozymes and a decrease in the number of hairpin ribozymes **(Fig. 3A-D)**. The competition dynamics was sensitive to temperature and Mg^2+^ concentrations **(Fig. 2E, 2F)**. Other factors studied also shifted the observed dynamics. For example, borate minerals have been suggested as a potential solution to the question of prebiotic synthesis of RNA by stabilizing ribose^5^. We found that three borate minerals previously studied in the context of ribose stabilization, colemanite (Ca_2_B_6_O_11_·5H_2_O), ulexite (NaCaB_5_O_6_(OH)_6_·5H_2_O), and kernite (Na_2_B_4_O_6_(OH)_2_·3H_2_O), showed differential effects on the activity of hammerhead and hairpin ribozymes **(Fig. S3)**.

**Figure 3.**
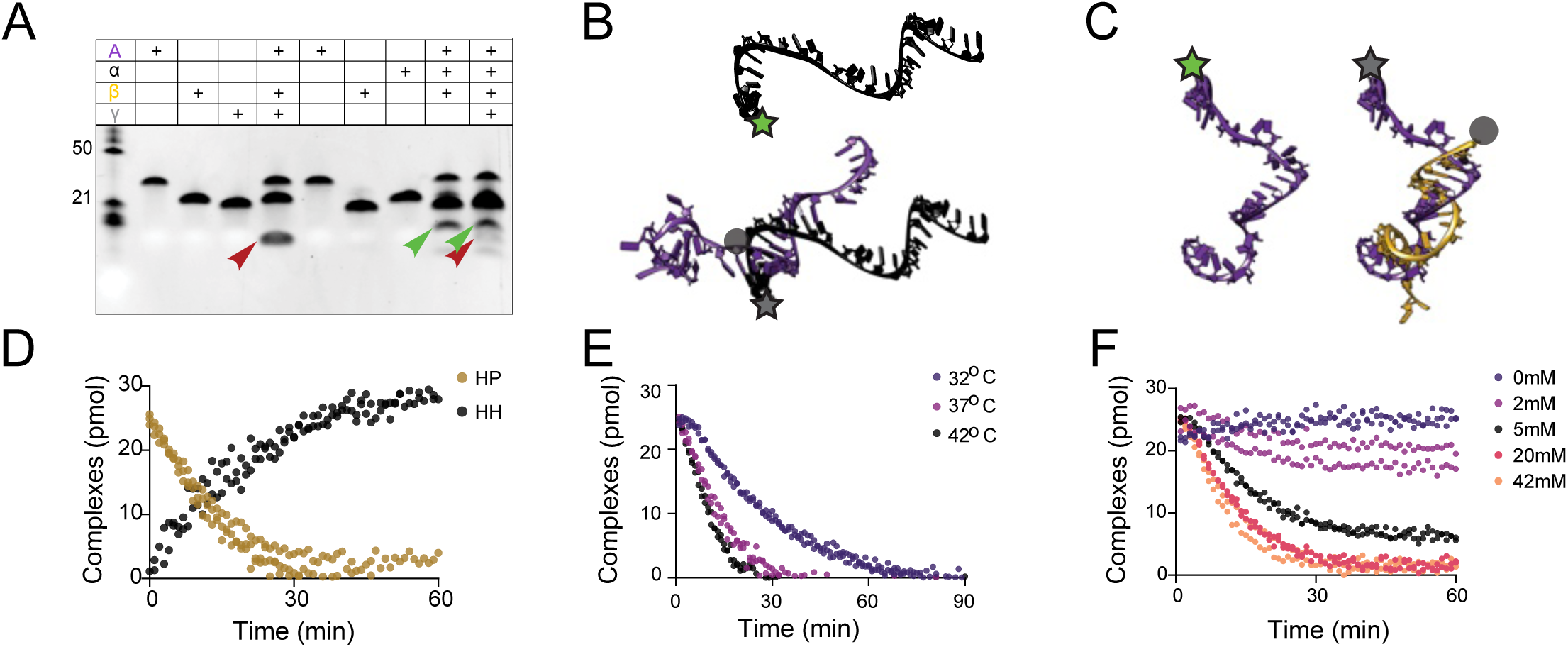
Analysis of the food chain functional dynamics. **A,** Gel electrophoresis results showing cleavage activity. Lane 5 (HP), Lane 9 (HH), and Lane 10 (food chain reaction) are highlighted, with arrows indicating the cleavage products. **B,** 3D model illustrates the hybridization between the α strand (black) with an attached fluorophore (star symbol) and the ancestral strand (purple) linked to a quencher (gray circle), leading to fluorescence quenching. **C,** Depicts the association of the ancestral strand (purple) with a fluorophore (star symbol) to the β strand (gold) that is connected to a quencher (gray circle), resulting in signal quenching. **D,** Graph showing the increase in HH and decrease in HP levels upon addition of the α strand. **E,** The graph displays the temperature’s effect on the system. Elevated temperatures expedite the degradation of HP by HH, **F,** The graph demonstrates the essential role of Mg++ ions in the food chain reaction, including a dose-response effect where increased Mg++ concentrations lead to enhanced HP degradation..

## Discussion

In this short study we reconstructed a minimal heterotrophic “food chain” in the RNA world by engineering hammerhead and hairpin ribozymes that both depend on a shared, Dead Sea–derived ancestral strand as a limiting resource. Using environmental sequences that are stable in hypersaline Dead Sea conditions, we showed that a single ancestral RNA can be partitioned into two distinct ribozyme species that compete over this strand, with one (hammerhead) acting as a predator that degrades the scaffold of the other (hairpin) to free additional ancestral copies and thereby promote its own assembly. The balance between these two “species” is tunable by environmental parameters such as temperature, Mg²⁺ concentration, and the presence of specific minerals, highlighting how small changes in prebiotic geochemistry could have reshaped early molecular ecologies. Our results suggest that food-chain-like interactions between simple RNA catalysts are chemically plausible and may have arisen naturally in geochemical niches similar to those sampled in the Dead Sea ^44^.

One feature of the design presented here is that the predator breaks down the prey to release a component which it then utilizes unchanged. This is reminiscent of modern ecosystems, where predators rarely exploit every part of their prey; for example, an animal may tear through skin and connective tissue primarily to access nutrient-rich organs or muscle. By analogy, our hammerhead ribozyme does not need to “use” every fragment generated from the prey hairpin complex. Its essential function is to cleave the hairpin scaffold in a way that liberates the limiting ancestral strand, thereby increasing the pool of material available for assembling additional hammerhead ribozymes. This dissociation between what is broken and what is ultimately incorporated suggests that early molecular predators could have been selected not for maximal recycling of all components, but for their efficiency in releasing just those parts needed to sustain their own replication.

It is interesting to note that the ancestral strand that seeds both ribozyme species is enriched in G, A, and C, with a marked paucity of U residues. This composition is consistent with Dead Sea stability data identifying G- and C-rich k-mers among the most stable sequences in hypersaline brines and resonates with hypotheses that early genetic systems may have been GC-biased or operated with a reduced alphabet ^45^.

We were not able to identify the source of RNA sequences used in this study. The Dead Sea system has complex geology and biology, most of it yet unmapped. Underwater springs in the Dead Sea originate from different sources ^44^. In the Samar/Darga area the springs derive from the Judean Mountain aquifer, whereas springs more to the south (e.g. Qedem) originate from deep thermal, hypersaline aquifers ^44^. Though recently described, these springs have been likely feeding the Dead Sea for a much longer period, entering the lake through tectonic faults that act as conduits. In the lake, these manifest as conical funnels in the sediment as well as holes in the salt layers accumulated on the bottom of the lake. The springs sampled in this study were located in the Samar/Darga area. The water of these springs is anoxic and sulfidic with a salinity range between fresh water (specific gravity 0.98 g cm^-3^) and sea-water like (specific gravity 1.029 g cm^-3^). Based on rare earth element analysis, this variation in salinity, as well as variation in other major ions was shown to be the result of different flow velocities and residence time in the underground before emerging into the Dead Sea. It was hypothesized that the major reason for the enhanced abundance of life around these springs is the formation of a microlayer with reduced salinity, however, these springs fluctuate randomly in flow intensity ^43,44^.

The food-chain design also points to a conceptual bridge between prebiotic heterotrophic RNA and modern therapeutic concepts based on nucleic acids. The hammerhead predator we describe is, in essence, a modular RNA catalyst that recognizes and dismantles another structured RNA while sparing a third strand it depends on. This arrangement is analogous to an extremely potent, self-amplifying “drug”: once a small pool of predator ribozymes is present, each catalytic turnover not only destroys target RNA (the prey scaffold) but also releases additional strands that can fuel the assembly of more predators. This “drug” would also be self-contained, decaying once all of its target has been destroyed.

Biology is traditionally framed around the cell as the basic unit of life, which can obscure molecular-scale ecologies that exist outside cellular envelopes. The interactions we reconstruct here involve entirely acellular players: short RNA strands that assemble into transient ribozymes, thought of here as actual organisms, compete for limiting resources, and engage in predator–prey-like relationships. These dynamics demonstrate that key ecological concepts—competition, predation, resource limitation, and niche specialization—can be instantiated at the level of naked RNA molecules in solution.

Taken together, our results support a view of the RNA world in which resource competition, predation, and environmental heterogeneity could arise even in small, compositionally simple systems. They also raise the fascinating hypothesis that heterotrophic RNA is a phenomenon that exists even today in some overlooked environments and niches, where transient consortia of RNA molecules - once a dominating life form on earth - assemble, compete, and decay in the shadow of cellular life.

## Methods

### Oligonucleotides

RNA oligonucleotides, including any modification, were ordered from Integrated DNA Technologies (IDT), reconstituted to 100 μM with ultrapure DNase/RNase free water (Biological Industries, Israel). RNA sequences are listed below.

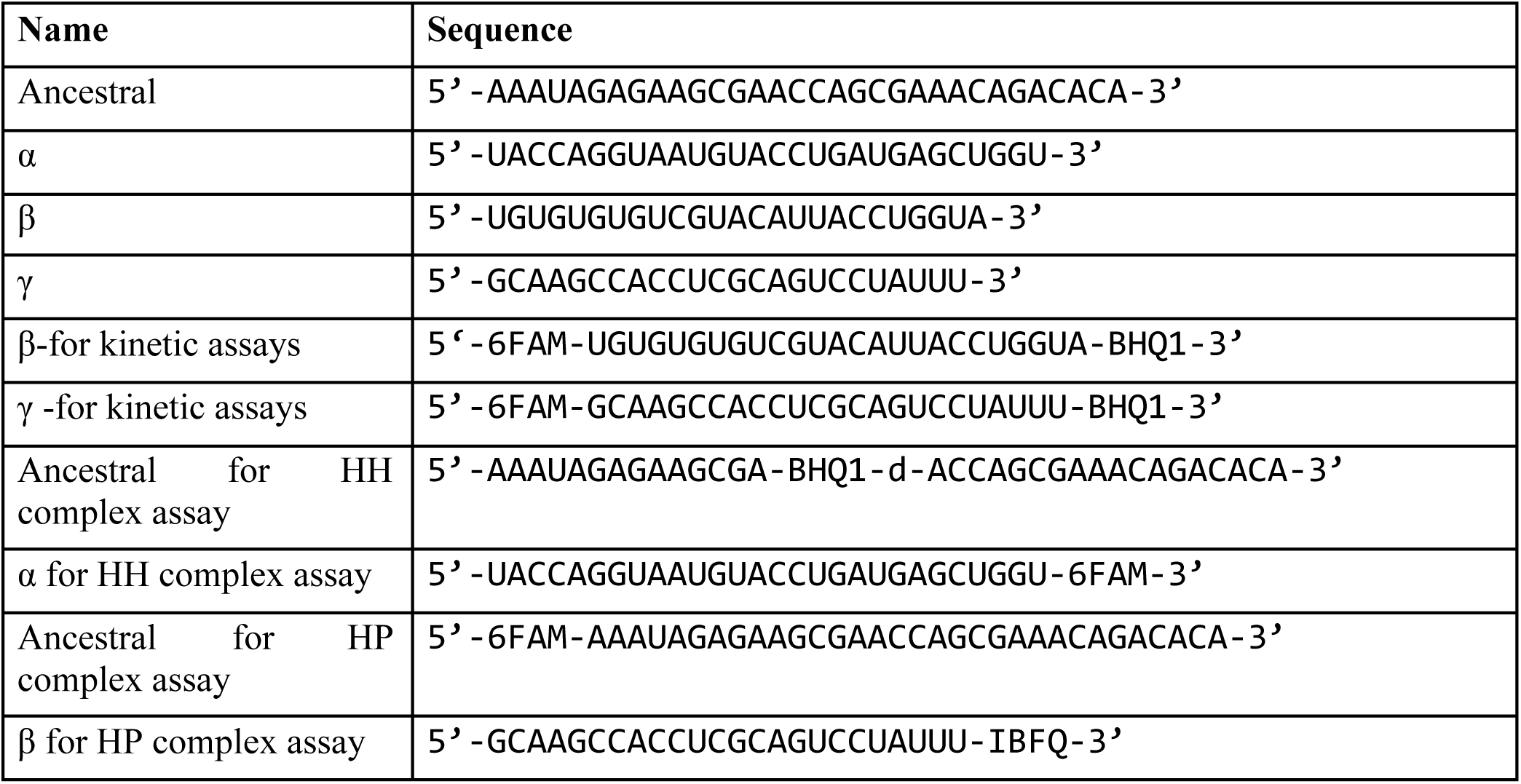

FAM- Fluorescein

BHQ1- Black hole 1 quencher

IBFQ- Iowa black FQ

Dead Sea RNA sequences are listed below.

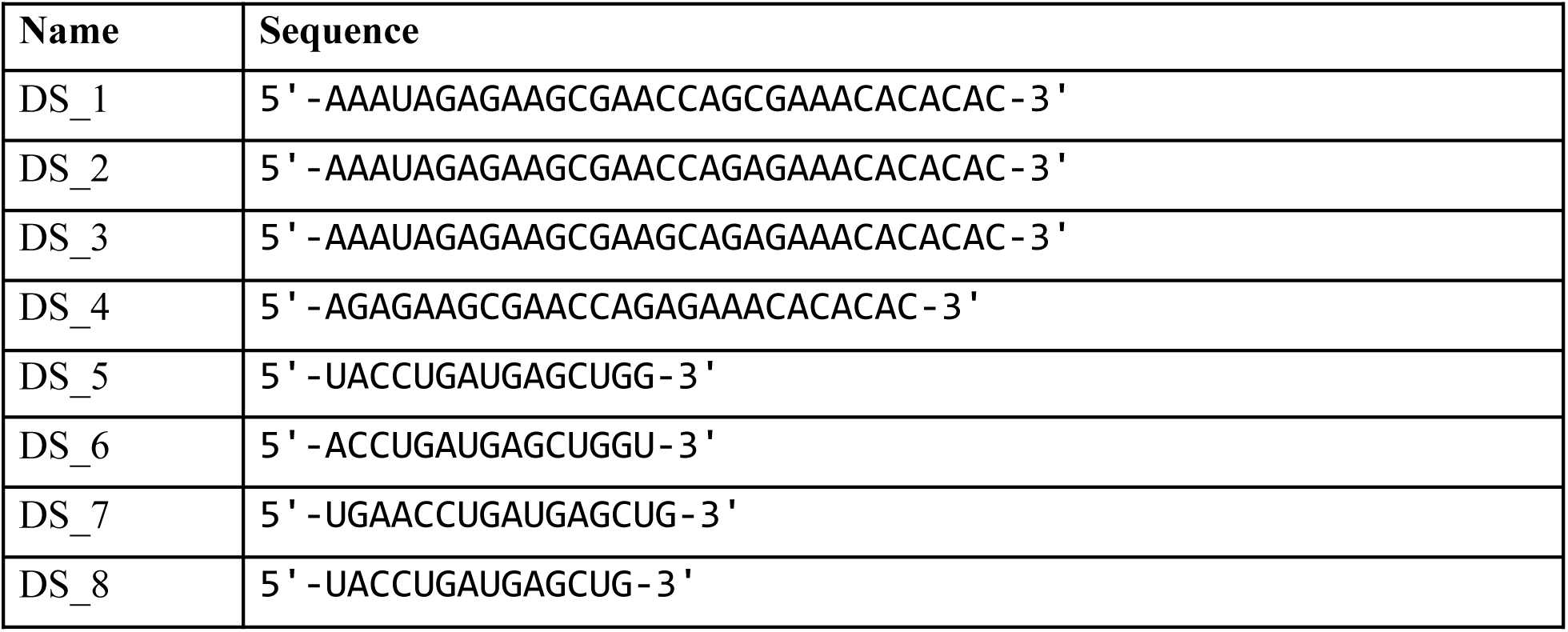
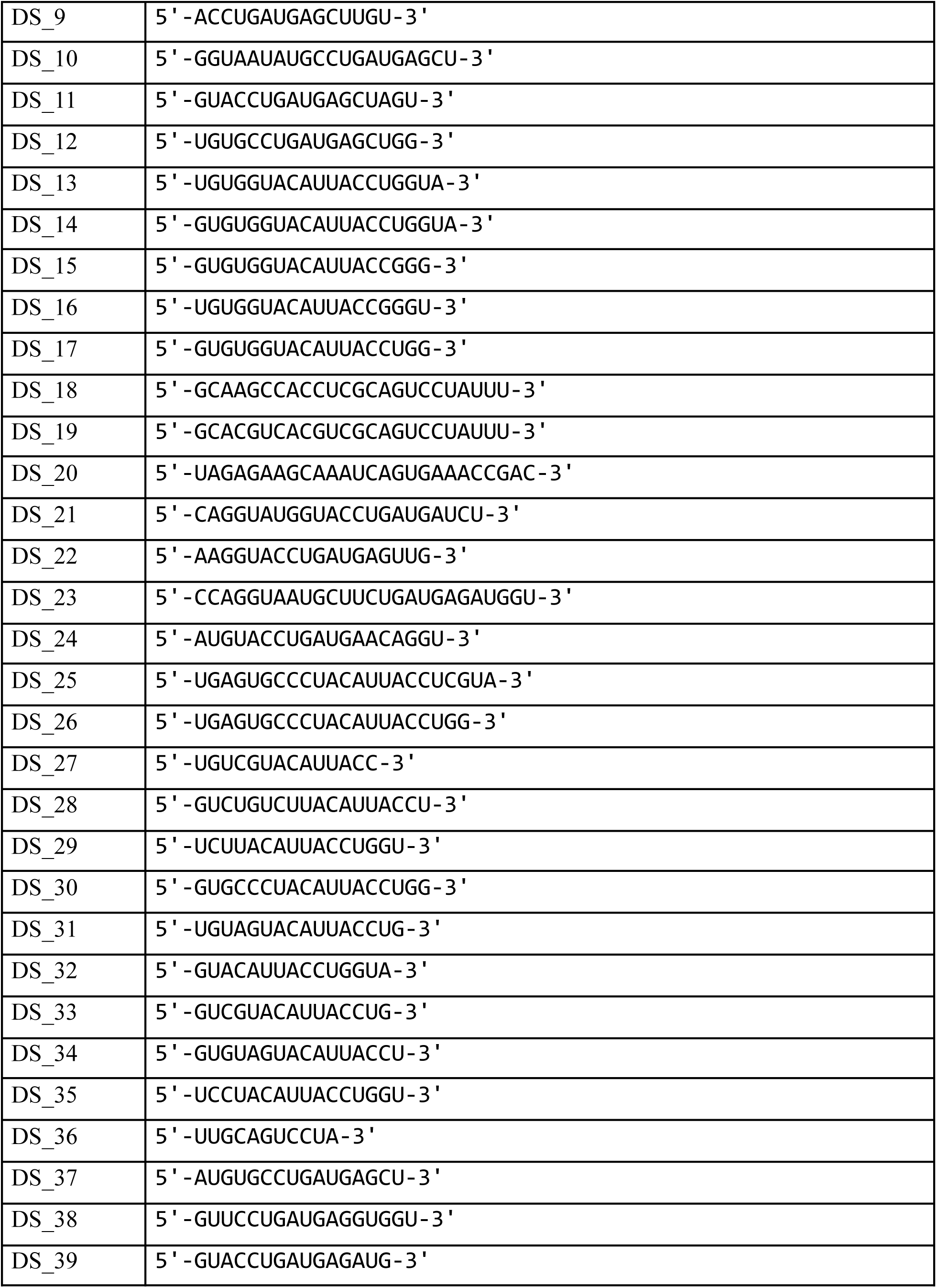
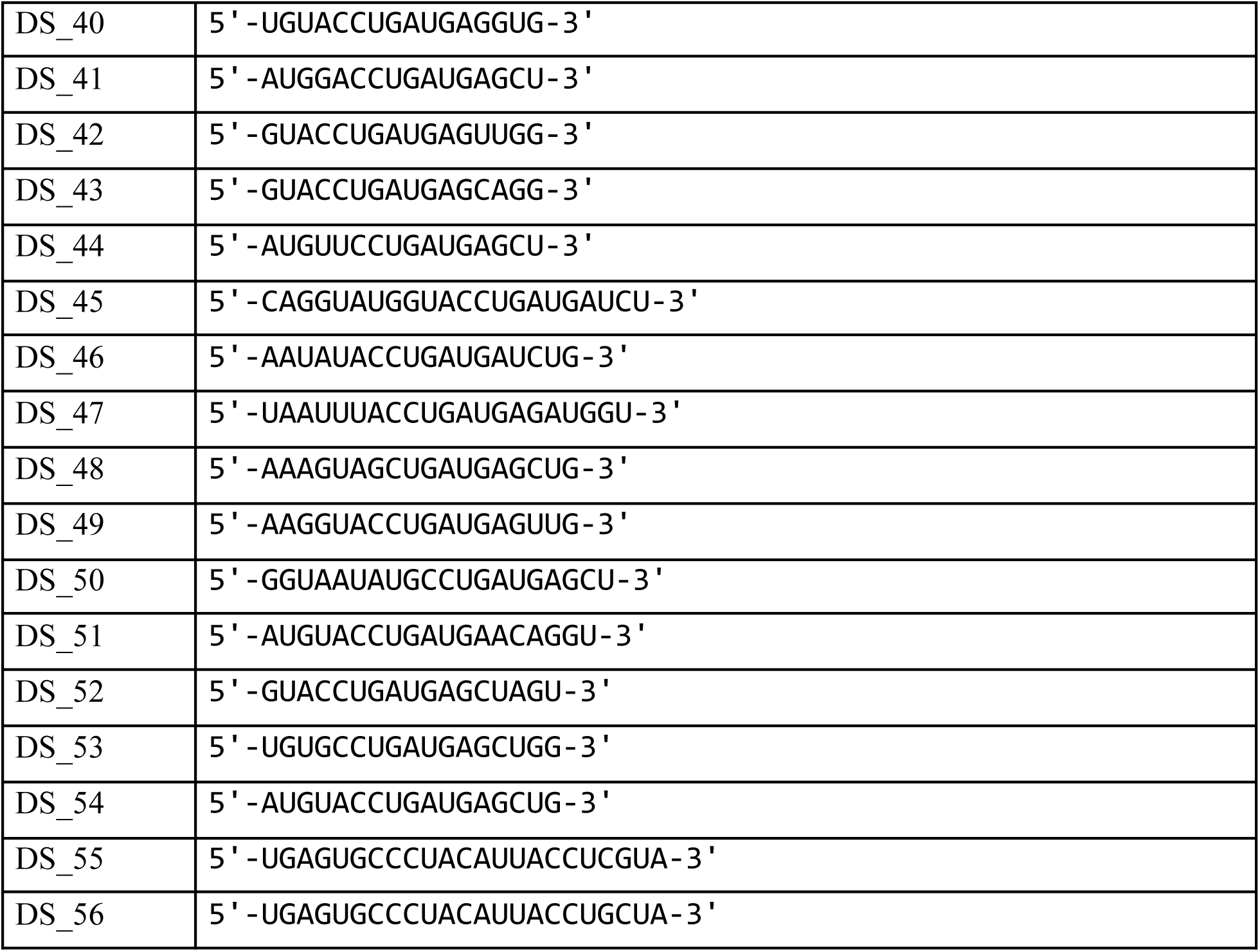

### Ribozyme activity assays

All components (two ribozyme strands and substrate) were mixed at a 1:1:1 ratio in Tris-acetate-EDTA (TAE) buffer (biological Industries, Israel) supplemented with 5 mM MgCl_2_ (Invitrogen, Catalog # AM9530G) or Mg(NO_3_)_2_ (Sigma-Aldrich, Catalog # 63084) at 37° C for 2 hrs in a Bio-Rad C1000 Touch Thermal Cycler. Final concentration of each component was 1 μM in 30 μL. Cleavage was verified by gel electrophoresis or Matrix-Assisted Laser Desorption/Ionization Mass Spectrometry (MALDI-MS) as described below.

### MALDI-MS

Matrix solution consisting of 12 mg/mL 2,4,6-trihydroxyacetophenone, 6 mg/mL 2,3,4-trihydroxyacetophenone, 5 mg/mL dihydrogen ammonium citrate was prepared in 50/50 acetonitrile/water. 0.5 µL of oligonucleotide solution and 0.5 µL matrix solution were mixed on the stainless steel MALDI plate and allowed to dry at atmospheric conditions. Samples were analyzed on a MALDI 8030 TOF instrument (Shimadzu Europa GmbH), operating in linear positive mode, allowing for the detection of molecules within a mass range of 2000–15000 Da.

### Gel electrophoresis

For RNA preparation, each sample was mixed with a 2X concentration of RNA loading dye (New England Biolabs, Catalog # B0363S). This mixture was subsequently incubated at 70°C for 10 min to facilitate denaturation. The denatured samples were then subjected to electrophoretic separation on 10% polyacrylamide Tris-borate-EDTA (TBE)-Urea gels (Invitrogen, Catalog # EC68752BOX). The electrophoresis was conducted in TBE buffer (Biological Industries, Israel) at a constant voltage of 180V for approximately 30 min. Post-separation, the gels were stained using SYBR™ Gold Nucleic Acid Gel Stain (Invitrogen, Catalog # S11494) for 10 min to visualize the RNA bands. For molecular size determination, both microRNA Marker (New England Biolabs, Catalog # N2102S) and Low Range ssRNA Ladder (New England Biolabs, Catalog # N0364S) were employed as ladder standards. Gel imaging was performed using a Cytiva Amersham ImageQuant 600 system to document the results.

### Kinetics assays

Measurement of the amount of complexes was performed based on a fluorophore-quencher assay. To assess the number of HH complexes, the α strand was synthesized with a 6-FAM at the 3’ and the ancestral strand was synthesized with a BHQ-1.The hybridization of the two strands (formation of the HH complex) resulted in quenching of the fluorescent signal. The quenching of the signal was in correlation to the number of complexes formed in the reaction. To assess the number of HP complexes, a similar technique was employed, but this time the ancestral strand was synthesized with 6-FAM at the 5’ and the β strand was synthesized with IBFQ quencher at the 3’. To calculate the amount of HP complexes from the fluorescent signal the following transformation was performed: The fluorescent signal from 6-FAM ancestral strand alone was considered as %0 complexes (*f*). The quenched signal (*q*) which resulted from mixing the 6-FAM ancestral strand with β strand conjugated to a quencher was considered as 100% complexes (25pmol). To calculate the pmol of complexes at each time point (*a*), the next formula was used:

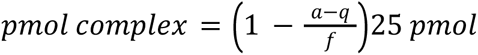

A similar method was performed to calculate the amount of HH complexes with a slight difference. The fluorescent signal was on the α strand and the quencher was on the ancestral strand. To assess the effect of temperature on the food chain reaction, the HH complex (described above) was set up as follows: 0.5 µM of the ancestral leg and β strand were mixed in 50 µl of TAE supplemented with 10 mM MgCl. After 1 hr of incubation (to allow hybridization and quenching) 0.5µM of the α strand was added. The addition of the strand resulted in the cleavage of the β strand and an increase in the fluorescent signal Reactions were setup in a 96-well plate suited for fluorescent measurement (Greiner, Catalog # 655090). Fluorescent measurement (495nm excitation and 530nm emission) was detected with Tecan infinite M plex, every minute for a duration of 180 min (60 min for hybridization + 120 min for cleavage reaction). The experiment was performed 3 times, each time with a different incubation temperature: 32°C, 37°C or 42 °C.

To test the effect of MgCl_2_ on the food chain reaction a similar experiment was performed, however the temperature was constant at 37^0^C, and various concentrations of MgCl_2_ were tested: 0, 2 mM, 5 mM, 20 mM or 42mM.

### RNA extraction

RNA extraction was performed on-site as quickly as possible from sampling. One liter of the sample was first passed through a 0.22 μm cellulose (Corning, Catalog # 430769) membrane using a vacuum-based filtration system. Subsequently, the flow-through was further filtered using a 0.22 μm nylon filter (Corning, Catalog # 430771). This nylon filter, containing retained negative particulates, was immediately frozen on dry ice for preservation. In the laboratory, the frozen membranes were shredded in a tube with scissors. To this, a mixture of 200 μL of Ultra-Pure Water (UPW), 400 μL of binding buffer from the Zymo RNA Extraction Kit (Zymo research, RNA Clean & Concentrator-25, Catalog # R1017) and 1.2mL of 100% Ethanol ( Biolab, Catalog # 525052100) was added. The tube was then vigorously shaken for 5 min to ensure thorough mixing. The contents were then transferred to a Zymo Column for RNA extraction, following the manufacturer’s protocol, which included a DNase (Invitrogen, Catalog # AM2238) treatment step to remove any contaminating DNA. The extracted RNA was stored at -80°C until further processing for sequencing.

### Library preparation and sequencing

8 μL from each RNA extraction sample were used for RNA library preparation with the D-Plex Small RNA-seq Kit (Diagenode, Catalog # C05030001). Indexes were attached using the D-Plex Unique Dual Indexes for Illumina Set A (Diagenode, Catalog # C05030021) and Set B (Diagenode, Catalog # C05030022) according to the manufacturer’s protocol. Library yield and size were measured by Agilent 4200 Tape-Station system, using a High Sensitivity D1000 ScreenTape (Agilent, Catalog # 5067-5584) or D1000 ScreenTape (Agilent, Catalog # 5067-5582). In addition, concentrations were measured with a Qubit 3 fluorometer System, using the Qubit® dsDNA HS Assay Kit (Invitrogen, Catalog # q32854). Sample multiplexing was prepared for 4 nM by diluting and pooling all samples equally. The pooled sample was sequenced on a NextSeq500 system (Illumina), using the NextSeq 500/550 High Output Kit v2.5, 75 Cycles (Illumina, Catalog # 20024906) or 150 Cycles (Illumina, Catalog # 20024907).

### Sequencing analysis

Bioinformatic analyses were performed using a custom in-house pipeline. Initially, reads were demultiplexed by Local Run Manager software (Illumina) and FASTQ files from distinct lanes merged for each sample. Subsequent read trimming was conducted with cutadapt (v4.4) ^46^ to remove UMIs, adapters and concatmers, and to maintain a Q-Score threshold of 30. In the metagenomics analysis phase, following the Kraken protocol ^47^, processed reads underwent classification via Kraken2 (v2.1.3 -) ^48^ against a RefSeq database of archaea, bacteria, viral, plasmid, human, protozoa, fungi and plant (as of 10/09/2023), with a k-mer size of 35. Reads corresponding to species detected in control samples were excluded from Dead Sea sample analysis. Classified reads served as the basis for taxon abundance estimates using Bracken ^49^. Visualization of hierarchical taxonomy, including Sankey and Krona plots, was achieved through KrakenTools, Pavian ^50^, and Krona (v2.8.1) ^51^. It should be noted that the kraken tools have a requirement of a minimum length of 35 nt for classification. Since a portion of the reads were shorter than 35 nt, we can assume the tool missed some known sequences.

### Inductively Coupled Plasma (ICP), pH and temperature measurement of samples

Ion measurement was performed by ICP. Both ICP and pH measurements were performed by Aminolab Ltd (Rehovot, Israel). A hierarchically clustered heatmap was generated, displaying the Z-score across each ion’s molarity. Z-score = (x - mean)/std. Temperature was measured on site with a waterproof probe test thermometer (Therma-cert, Catalog #TC-004).

### RNA stability assay

Three RNA libraries of 18N were ordered (IDT) at quantities enabling the creation of dozens of microliters in volume of 1mM library. The total number of different sequences coded by an 18N library is 4^18^=∼6.87*10^10^. Considering the total number of sequences in a representative library we created (∼2*10^16^), ∼2.8*105 copies represent each unique sequence. Each library was diluted at 1:10 volume ratio with DDW or Dead Sea water (5µ 18N library + 45µl DDW/Dead Sea water in a 200µl PCR tube) and incubated at 30°C for 7 days with Bio-Rad C1000 Touch Thermal Cycler. The procedure repeated two additional times creating triplicates from each library. Samples were taken for sequencing at t=0 and t=7 days from the two libraries, overall 9 samples. As a control, we used a known RNA sequence, 18 nucleotides long and diluted it to the same concentrations of the RNA libraries dilution, same incubation conditions as well. Post-processing, sequences were filtered to retain only 18-nucleotide (nt) long reads. Furthermore, the K-mers (K=6) of each sample were counted, and their proportion was calculated for normalization. A hierarchically clustered heatmap was generated based on the frequency of each K-mer in every sample. The K-mers frequencies were treated as RNA-Seq count data. For each K-mer, the following values were calculated: 1 Log_2_ Fold Change, derived from the log-transformed ratio of mean frequencies between t=0 and t=14; 2) P-value, from a Student’s t-test comparing frequencies at t=0 and t=14 (n=3), with subsequent False Discovery Rate (FDR) adjustment for multiple comparisons. Using these parameters, a Volcano plot was generated, highlighting K-mers with notable stability variations in Dead Sea water. We sought ancestral K-mers (K=3,4,5,6) among those significantly altered, evaluating their presence in the Dead Sea samples. Ancestral strand coverage was quantified as the percentage covered by stable K-mers. For controls, coverage was similarly assessed for both a scrambled ancestral strand (matching base composition, 30 repeats) and a random 32 nt sequence (uniform base probability of 0.25, 30 repeats), employing one-sample t-tests to ascertain statistical significance.

### Ribozymes’ oligonucleotides in Dead Sea

Ancestral, α, and β strand sequences underwent Blast analysis (v2.14.1+)^52^ against processed Dead Sea reads, employing a word size of 7 and an E-value threshold of 100. Selection criteria for hits mandated intact catalytic cores—defined according to the literature ^37,53–55^, excluding those with gaps, misalignments, or originating from negative strands. A scoring system was devised to identify optimal hits for active ribozyme construction. In this system, nucleotides aligned with their counterparts in the reference sequences (ancestral, α, or β) contribute to the score as 1/(index distance from the catalytic core), enhancing the selection of sequences with proximity to essential catalytic regions. Conversely, mismatches contribute a score of 0, inherently prioritizing matches closer to the ribozyme’s functional core and thus promoting effective hybridization and functional integrity. Further refinement in hit selection incorporated additional parameters: mismatches within catalytic cores, hit and hybridization lengths, and predicted 2D structures using Nupack (v4.0.0.28) ^56^. These criteria ensure the selection of sequences most likely to form active ribozymes, considering both sequence fidelity and structural compatibility. Candidate strands were ordered from IDT, and their functionality was tested utilizing Gel electrophoresis as described above.

### A phylogenetic tree of the ancestral strand in Dead Sea samples

Blast hits of the ancestral sequence, derived from Dead Sea samples sequencing data and exceeding 30 nucleotides from the positive strand, were selectively filtered. Subsequently, a phylogenetic tree of these ancestral sequences was constructed. The construction of the tree entailed a reference-based Multiple Sequence Alignment (MSA) using MAFFT (v7.520)^52^, tree construction with IQ-TREE (v2.0.7)^57^, and visualization employing ETE3 (v3.1.3)^58^.

### Assessment of ribozyme activity in Dead Sea samples

All components (two ribozyme strands and substrate) were mixed at a 1:1:1 ratio in one of the 3 Dead Sea samples at 37 °C for 2 hrs in a Bio-Rad C1000 Touch Thermal Cycler. The final concentration of each part was 1 μM in 30 μL. Desalting was performed with an RNA Extraction Kit (Zymo research, RNA Clean & Concentrator-25, Catalog # R1017) following the manufacturer’s protocol. Cleavage was verified by gel electrophoresis as described above.

### 3D modeling

Three-dimensional model prediction of ribozymes was conducted using Farfar2^59^, and visualized with Chimera^60^.

### The effect of Mg^++^ on the HH and HP

All components (two ribozyme strands and substrate) were mixed at a 1:1:1 ratio in TAE buffer (biological Industries, Israel) supplemented with 0,1, 2.5, 5 or 10 mM MgCl_2_ at 37 °C in a Bio-Rad C1000 Touch Thermal Cycler. The HP was incubated for 30 min and the HH was incubated for 2 hrs. Final concentration of each part was 1 μM in 30 μL. Cleavage was verified by gel electrophoresis as described above.

### Ribozyme kinetics assays - Direct

The kinetics of the ribozyme were measured using a fluorophore-quencher based assay. The substrates for the ribozymes were conjugated to a 6-Fam at the 5’ and a BHQ1 to the 3’. The cleavage of the substrate by the ribozyme disrupts the quenching interaction between the fluorophore and quencher, leading to the production of a detectable fluorescent signal. Reactions were set up in a 96-well plate suited for fluorescent measurement (Greiner, Catalog # 655090). Each well contained 50 µl of TAE buffer, 5mM MgCl_2_, ribozyme (HR or HP) and it’s substrate at a molar ratio of 1:5 in favor of the substrate (0.1µM ribozyme and 0.5 µM substrate) or in favor of the ribozyme (0.5µM ribozyme and 0.1 µM substrate). Fluorescent measurement (495 nm excitation and 530 emission) was detected with Tecan infinite M plex, every min for a duration of 180 min at 37 °C. To assess the effect of kernite, varying concentrations were used. Our stock solution of kernite was 140mM, corresponding to 4% w/v solution. Kernite mineral was purchased from “rock your chakra” and verified using ICP MS by Milouda laboratories, Kiryat Shmona, Israel). Emax, Vmax and K obs values (left, middle and right, respectively) calculated for both HPR and HHR at excess RZ and excess substrate conditions as follows: Maximum value (Emax) was the highest value of fluorescent signal throughout the whole experiment time. Maximum velocity (Vmax) was the slope of the linear curve extracted for the first 5 minutes of the reaction. K observed (Kobs) was calculated by the following equation: ln2/t_1/2_ (t_1/2_ is half time to reach Emax).

## Acknowledgements

The authors are grateful to the entire team at Augmanity, particularly Dr. R. Spokoini-Stern and Dr. Y. Amir, for their valuable discussions and technical expertise. We also extend our thanks to Dr. R. Ionescu for his assistance with Dead Sea sample collection, Dr. A Baumeister for his MALDI-TOF expertises and to G. Golan and A. Terkel for their contributions of Dead Sea imagery.

## Author contributions

The HP and HH food chain ribozymes were designed by IB, RW, and OBD. YS, NM, and RW carried out the experiments to assess the functionality of these ribozymes. Further investigation into the impact of kernite, magnesium ions (Mg++), and temperature on the food chain was conducted by SY and RW. Sample collection from the Dead Sea was accomplished by YS, NM, SY, RW, AI, and DI. The process of RNA extraction was undertaken by YS, NM, and RW, while library preparation and sequencing were executed by ALZ and NJ. CF and NV were responsible for the bioinformatic analysis. IB provided supervision for the entire research project and was the primary author of the manuscript.

## Supplementary figures

**Figure 1.**
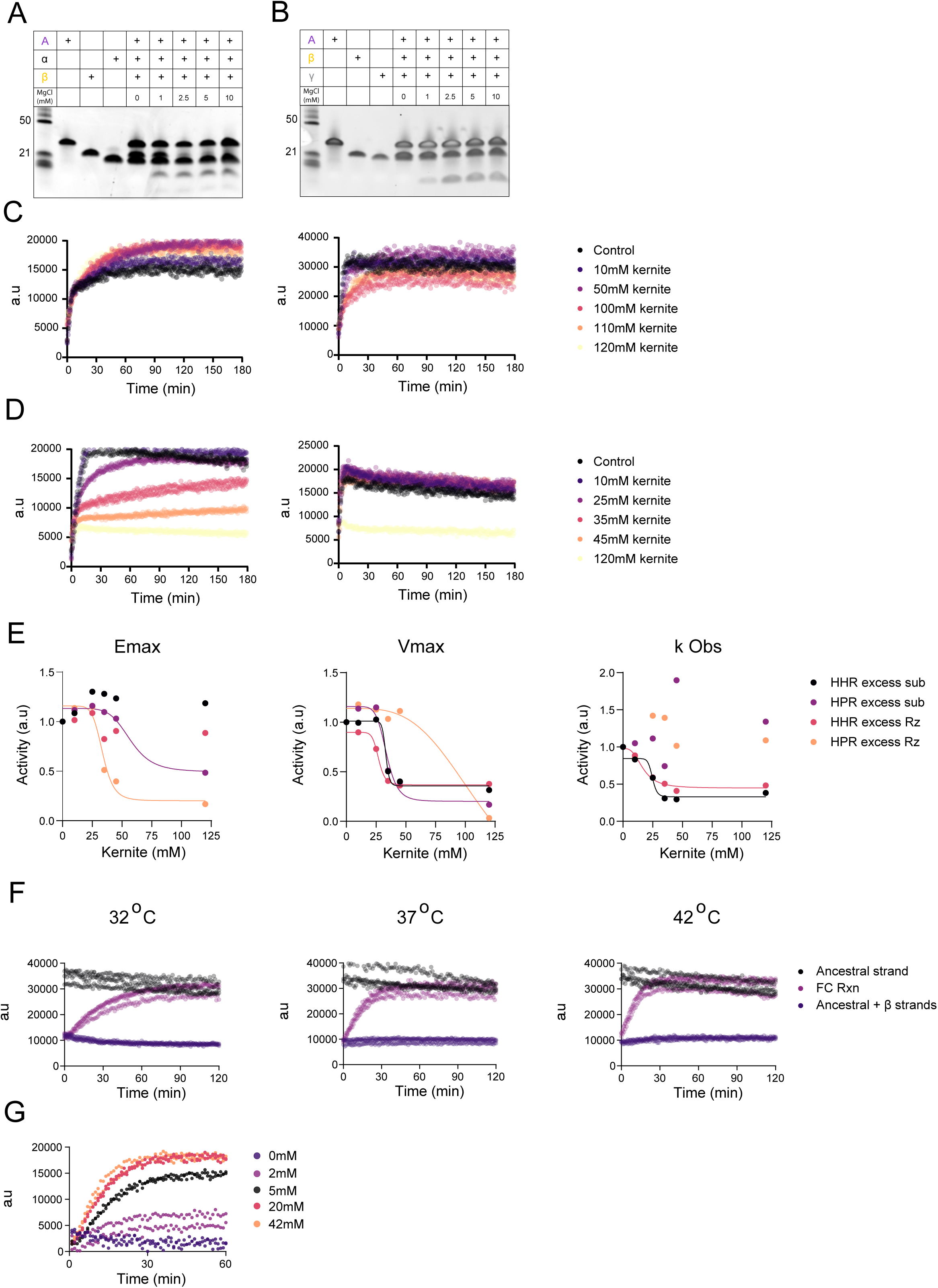
**A,** Gel electrophoresis demonstrating the effect of Mg^++^ on the activity of the HH (left gel) and HP (right gel) **B,** HHR kinetics at excess substrate or ribozyme (left and right respectively) at varying concentrations of kernite, ranging from none (control) to 120mM **C,** HPR kinetics at excess substrate or ribozyme (left and right respectively) at varying concentrations of kernite, ranging from none (control) to 120mM **D,** Emax, Vmax and K obs values (left, middle and right, respectively) calculated for both HPR and HHR at excess RZ and excess substrate conditions **E,** The graphs display the degradation of HP by HH in 3 different temperatures **F,** The graph demonstrates the essential role of Mg++ ions in the food chain reaction, including a dose-response effect where increased Mg++ concentrations lead to enhanced HP degradation.

**Figure 2.**
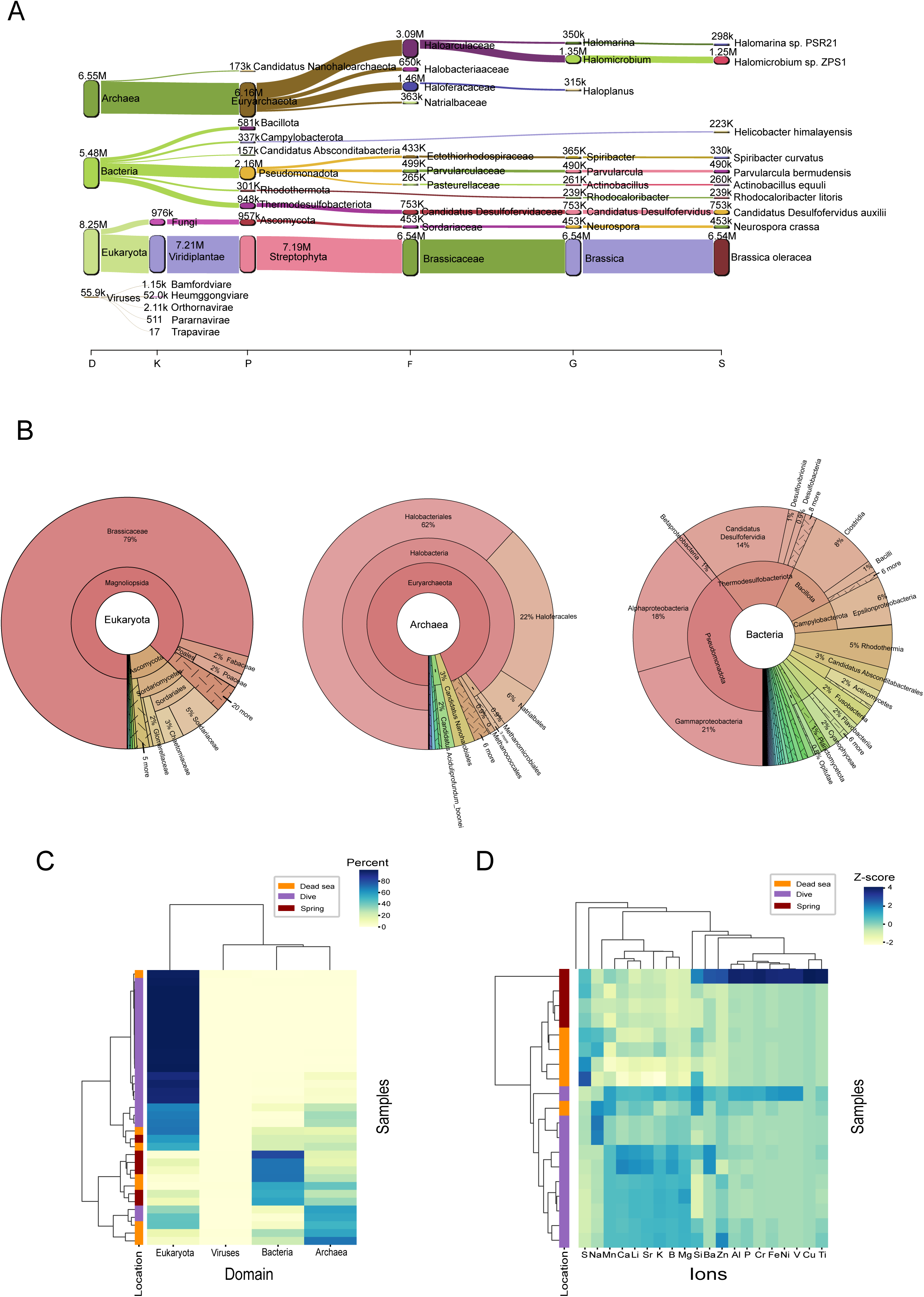
**A,** Sankey diagram of hierarchical taxonomic classification from Dead Sa taxonomic analysis. Nodes from left to right represent Domain, Kingdom, phylum, Family, Genus and Species. The links show the flow of taxonomic units between different taxonomic levels. **B,** Krona plots of Eukaryota, Ahrchaea, and Bacteria from Dead Sea metagenomic analysis. Each pie chart layer represents a taxonomic level, from inner to outer layer: phylum, Family, Genus and Species. **C,** Hierarchically clustered heatmap showing the proportion of species from each domain in each sample. Label bar on the left indicates the location from which samples were collected. **D,** A hierarchically clustered heatmap displaying the Z-score across each ion’s molarity. Z-score = (x - mean)/std.

**Figure 3.**
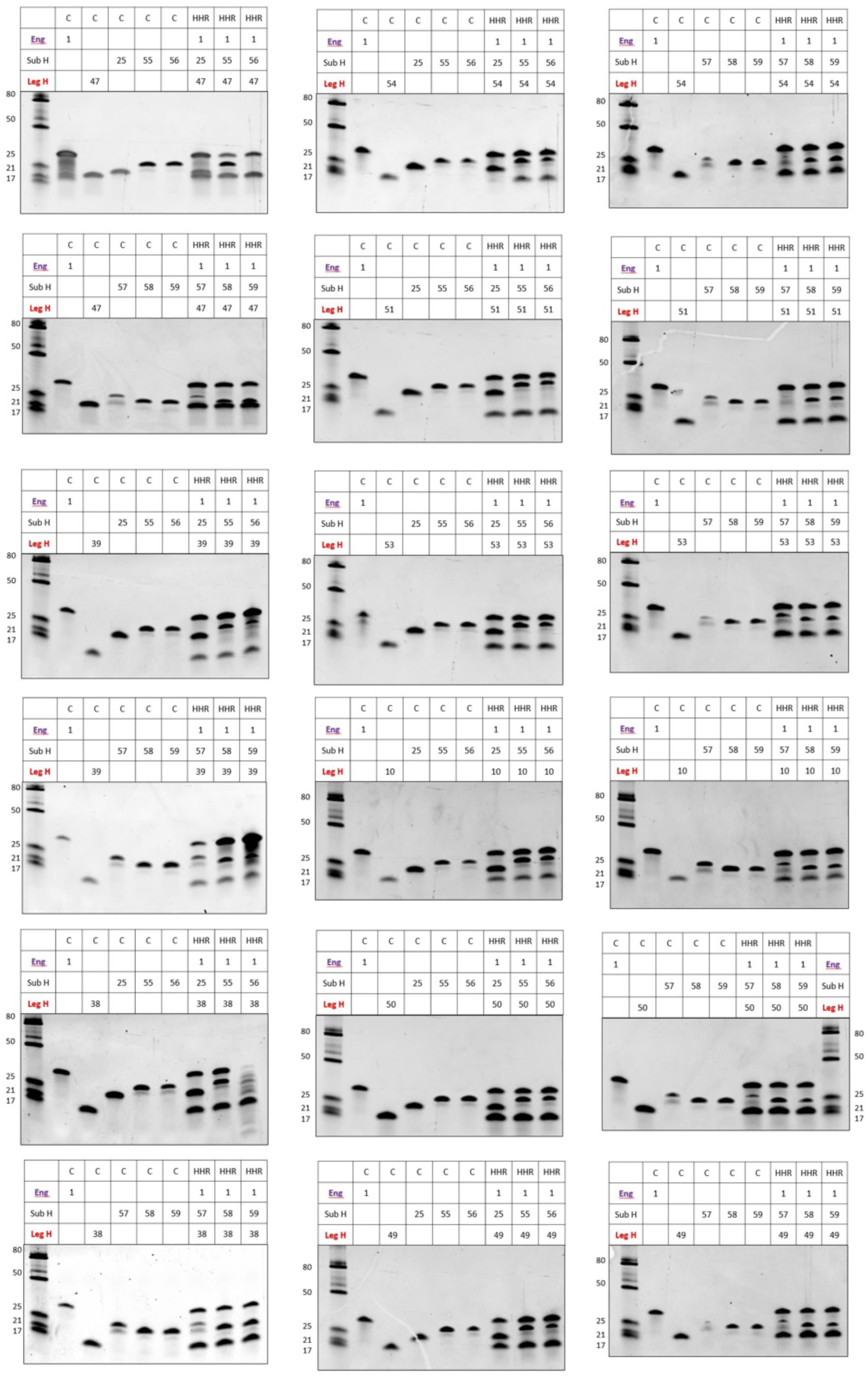
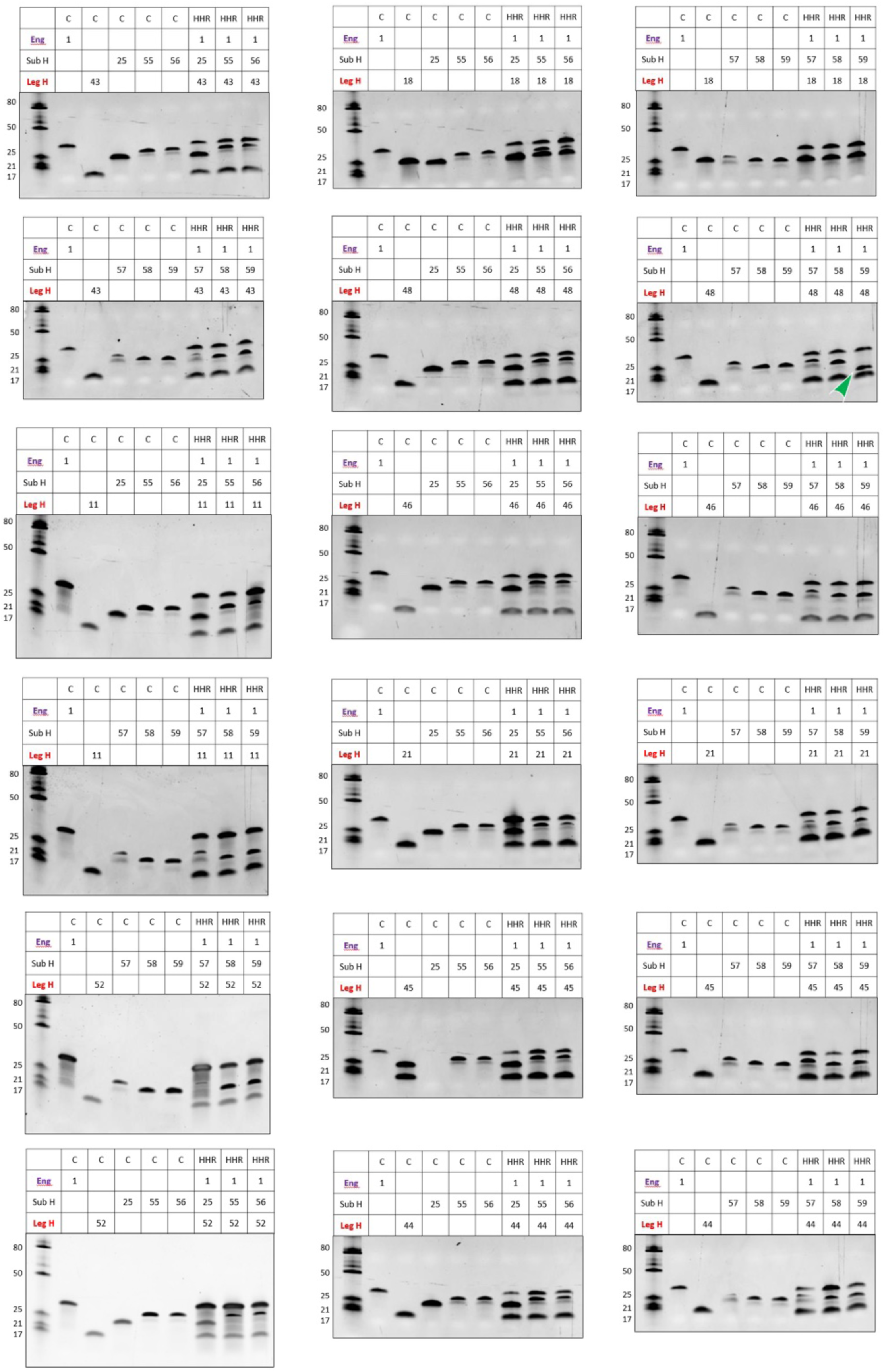
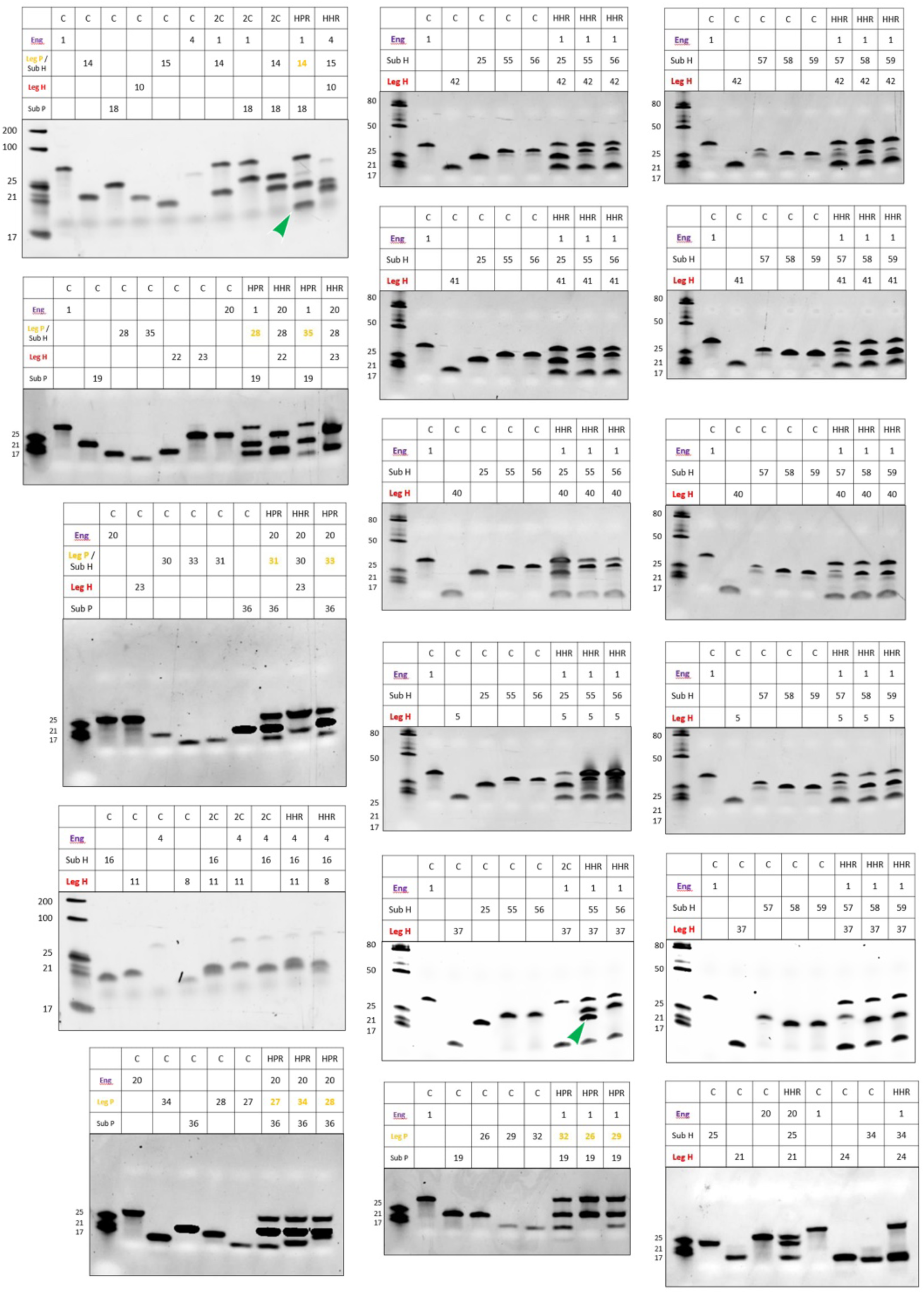
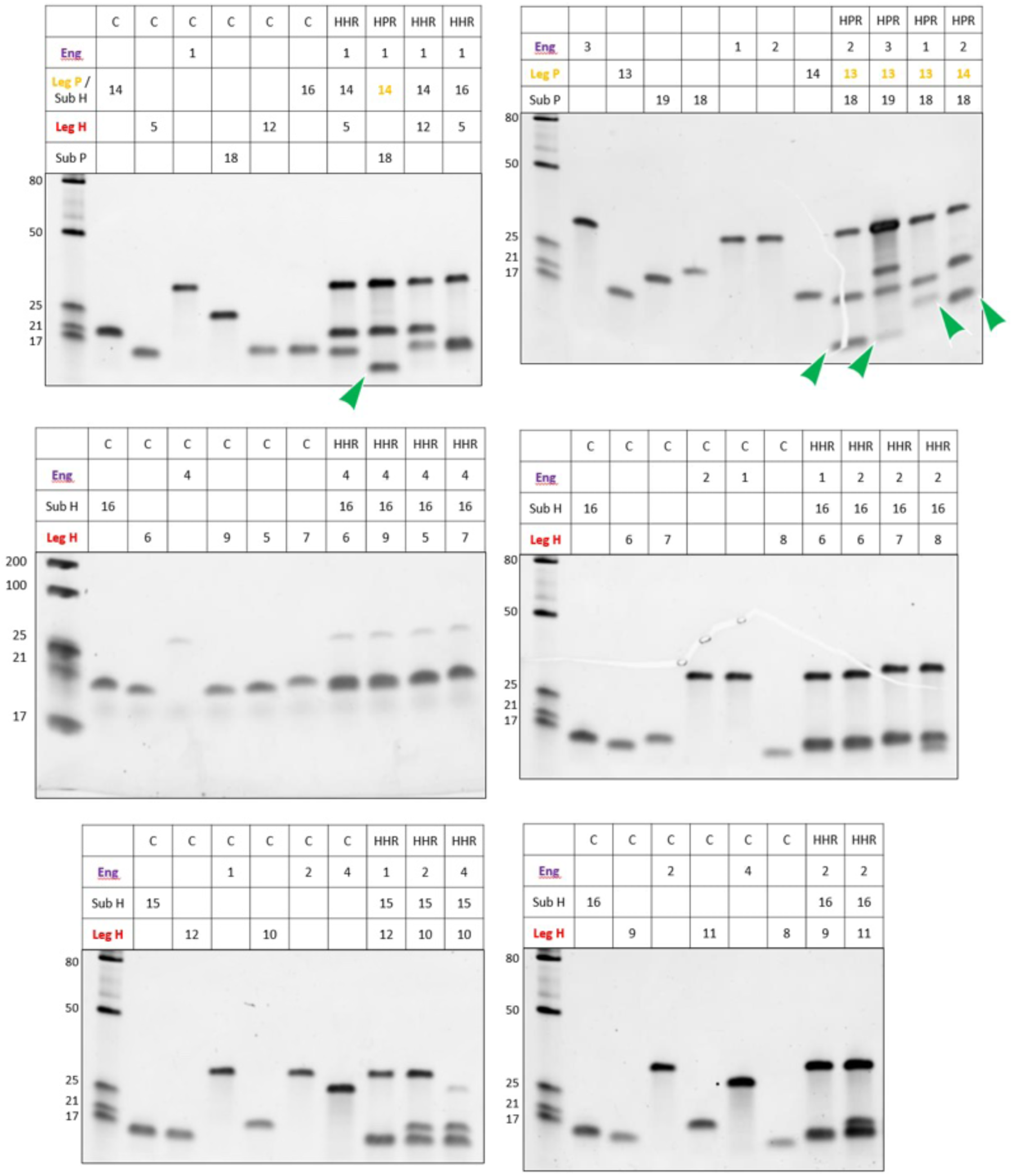
Gel Electrophoresis Analysis of Ribozyme Cleavage Activity. This figure displays the results of gel electrophoresis aimed at examining the cleavage activity of potential ribozymes isolated from the Dead Sea. “C” denotes the control lane, providing a baseline for comparison. “HHR” and “HPR” lanes represent the testing of potential HHR and HPR ribozymes, respectively. Arrows indicate the presence of cleavage products, highlighting the enzymatic activity of the tested ribozymes.

## References

1. Williams-Ashman, G. Horizons in Biochemistry. Albert Szent-Györgyi Dedicatory Volume Edited by Michael Kasha and Bernard Pullman. Perspectives in Biology and Medicine vol. 6 264–267 Preprint at 10.1353/pbm.1963.0024 (1963).

2. Neveu, M., Kim, H.-J. & Benner, S. A. The ‘strong’ RNA world hypothesis: fifty years old. Astrobiology 13, 391–403 (2013).

3. Gilbert, W. Origin of life: The RNA world. Nature vol. 319 618–618 Preprint at 10.1038/319618a0 (1986).

4. Rajamani, S. et al. Lipid-assisted synthesis of RNA-like polymers from mononucleotides. Orig. Life Evol. Biosph. 38, 57–74 (2008).

5. Ricardo, A., Carrigan, M. A., Olcott, A. N. & Benner, S. A. Borate minerals stabilize ribose. Science 303, 196 (2004).

6. Huang, W. & Ferris, J. P. One-Step, Regioselective Synthesis of up to 50-mers of RNA Oligomers by Montmorillonite Catalysis. Journal of the American Chemical Society vol. 128 8914–8919 Preprint at 10.1021/ja061782k (2006).

7. Swadling, J. B., Coveney, P. V. & Greenwell, H. C. Clay minerals mediate folding and regioselective interactions of RNA: a large-scale atomistic simulation study. J. Am. Chem. Soc. 132, 13750–13764 (2010).

8. Monnard, P.-A., Kanavarioti, A. & Deamer, D. W. Eutectic phase polymerization of activated ribonucleotide mixtures yields quasi-equimolar incorporation of purine and pyrimidine nucleobases. J. Am. Chem. Soc. 125, 13734–13740 (2003).

9. Schuster, G. B., Cafferty, B. J., Karunakaran, S. C. & Hud, N. V. Water-Soluble Supramolecular Polymers of Paired and Stacked Heterocycles: Assembly, Structure, Properties, and a Possible Path to Pre-RNA. Journal of the American Chemical Society vol. 143 9279–9296 Preprint at 10.1021/jacs.0c13081 (2021).

10. Cafferty, B. J., Fialho, D. M., Khanam, J., Krishnamurthy, R. & Hud, N. V. Spontaneous formation and base pairing of plausible prebiotic nucleotides in water. Nat. Commun. 7, 11328 (2016).

11. Hieronymus, R. & Müller, S. Towards Higher Complexity in the RNA World: Hairpin Ribozyme Supported RNA Recombination. ChemSystemsChem vol. 3 Preprint at 10.1002/syst.202100003 (2021).

12. Striggles, J. C., Martin, M. B. & Schmidt, F. J. Frequency of RNA-RNA interaction in a model of the RNA World. RNA 12, 353–359 (2006).

13. Tjhung, K. F., Shokhirev, M. N., Horning, D. P. & Joyce, G. F. An RNA polymerase ribozyme that synthesizes its own ancestor. Proc. Natl. Acad. Sci. U. S. A. 117, 2906–2913 (2020).

14. Horning, D. P. & Joyce, G. F. Amplification of RNA by an RNA polymerase ribozyme. Proc. Natl. Acad. Sci. U. S. A. 113, 9786–9791 (2016).

15. Jheeta, S. & Joshi, P. C. Prebiotic RNA synthesis by montmorillonite catalysis. Life 4, 318–330 (2014).

16. Attwater, J., Raguram, A., Morgunov, A. S., Gianni, E. & Holliger, P. Ribozyme-catalysed RNA synthesis using triplet building blocks. eLife vol. 7 Preprint at 10.7554/elife.35255 (2018).

17. Lincoln, T. A. Self-Sustained Replication of an RNA Enzyme. (ProQuest, 2009).

18. Wachowius, F. & Holliger, P. Non-Enzymatic Assembly of a Minimized RNA Polymerase Ribozyme. ChemSystemsChem vol. 1 12–15 Preprint at 10.1002/syst.201900004 (2019).

19. Mutschler, H., Wochner, A. & Holliger, P. Freeze-thaw cycles as drivers of complex ribozyme assembly. Nat. Chem. 7, 502–508 (2015).

20. Turk, R. M., Chumachenko, N. V. & Yarus, M. Multiple translational products from a five-nucleotide ribozyme. Proc. Natl. Acad. Sci. U. S. A. 107, 4585–4589 (2010).

21. Yarus, M. Ahead and behind: a small, small RNA world. RNA 21, 769–770 (2015).

22. Yarus, M. Darwinian Behavior in a Cold, Sporadically Fed Pool of Ribonucleotides. Astrobiology vol. 12 870–883 Preprint at 10.1089/ast.2012.0860 (2012).

23. Dawkins, R., Dawkins & Charles Simonyi Professor of the Public Understanding of Science Richard Dawkins. The Selfish Gene. (Oxford University Press, USA, 1976).

24. Weinberg, C. E., Weinberg, Z. & Hammann, C. Novel ribozymes: discovery, catalytic mechanisms, and the quest to understand biological function. Nucleic Acids Res. 47, 9480–9494 (2019).

25. Winkler, W. C., Nahvi, A., Roth, A., Collins, J. A. & Breaker, R. R. Control of gene expression by a natural metabolite-responsive ribozyme. Nature 428, 281–286 (2004).

26. Müller, S., Appel, B., Balke, D., Hieronymus, R. & Nübel, C. Thirty-five years of research into ribozymes and nucleic acid catalysis: where do we stand today? F1000Res. 5, (2016).

27. Hieronymus, R., Godehard, S. P., Balke, D. & Müller, S. Hairpin ribozyme mediated RNA recombination. Chem. Commun. 52, 4365–4368 (2016).

28. Forster, A. C. & Symons, R. H. Self-cleavage of virusoid RNA is performed by the proposed 55-nucleotide active site. Cell 50, 9–16 (1987).

29. de la Peña, M. & García-Robles, I. Intronic hammerhead ribozymes are ultraconserved in the human genome. EMBO Rep. 11, 711–716 (2010).

30. de la Peña, M. & García-Robles, I. Ubiquitous presence of the hammerhead ribozyme motif along the tree of life. RNA 16, 1943–1950 (2010).

31. Jimenez, R. M., Delwart, E. & Lupták, A. Structure-based search reveals hammerhead ribozymes in the human microbiome. J. Biol. Chem. 286, 7737–7743 (2011).

32. Perreault, J. et al. Identification of hammerhead ribozymes in all domains of life reveals novel structural variations. PLoS Comput. Biol. 7, e1002031 (2011).

33. Côté, F. & Perreault, J. P. Peach latent mosaic viroid is locked by a 2’,5’-phosphodiester bond produced by in vitro self-ligation. J. Mol. Biol. 273, 533–543 (1997).

34. Uhlenbeck, O. C. A small catalytic oligoribonucleotide. Nature 328, 596–600 (1987).

35. Kharma, N. et al. Automated design of hammerhead ribozymes and validation by targeting the PABPN1 gene transcript. Nucleic Acids Res. 44, e39 (2016).

36. Peña, M. de la, de la Peña, M., García-Robles, I. & Cervera, A. The Hammerhead Ribozyme: A Long History for a Short RNA. Molecules vol. 22 78 Preprint at 10.3390/molecules22010078 (2017).

37. Hieronymus, R. & Müller, S. Engineering of hairpin ribozyme variants for RNA recombination and splicing. Ann. N. Y. Acad. Sci. 1447, 135–143 (2019).

38. Shoval, S. & Zlatkin, O. Climatic changes during the Pliocene as observed from climate-sensitive rocks and clay minerals of the Sedom formation, the Dead Sea Basin. Clay Miner. 44, 469–486 (2009).

39. Changes in the thermo-haline structure of the Dead Sea: 1979–1984. Earth Planet. Sci. Lett. 84, 109–121 (1987).

40. Oren, A. The dying Dead Sea: The microbiology of an increasingly extreme environment. Lakes Reserv.: Res. Manage. 15, 215–222 (2010).

41. Wilkansky, B. Life in the Dead Sea. Nature 138, 467–467 (1936).

42. Website. Oren, A., Gurevich, P., Anati, D.A., Barkan, E., Luz, B. (1995) A bloom of Dunaliella parva in the Dead Sea in 1992: biological and biogeochemical aspects. Hydrobiologia 297, 173–185 10.1007/BF00019283.

43. Häusler, S., Weber, M., de Beer, D. & Ionescu, D. Spatial distribution of diatom and cyanobacterial mats in the Dead Sea is determined by response to rapid salinity fluctuations. Extremophiles 18, 1085–1094 (2014).

44. Ionescu, D. et al. Microbial and chemical characterization of underwater fresh water springs in the Dead Sea. PLoS One 7, e38319 (2012).

45. Higgs, P. G. & Lehman, N. The RNA World: molecular cooperation at the origins of life. Nat Rev Genet 16, 7–17 (2015).

46. Martin, M. Cutadapt removes adapter sequences from high-throughput sequencing reads. EMBnet.journal 17, 10–12 (2011).

47. Lu, J. et al. Metagenome analysis using the Kraken software suite. Nature Protocols 17, 2815–2839 (2022).

48. Wood, D. E., Lu, J. & Langmead, B. Improved metagenomic analysis with Kraken 2. Genome Biology 20, 257 (2019).

49. Lu, J., Breitwieser, F. P., Thielen, P. & Salzberg, S. L. Bracken: estimating species abundance in metagenomics data. PeerJ Comput. Sci. 3, e104 (2017).

50. Website. https://doi.org/10.1093/bioinformatics/btz715 doi:10.1093/bioinformatics/btz715.

51. Ondov, B. D., Bergman, N. H. & Phillippy, A. M. Interactive metagenomic visualization in a Web browser. BMC Bioinformatics 12, 385 (2011).

52. Altschul, S. F., Gish, W., Miller, W., Myers, E. W. & Lipman, D. J. Basic local alignment search tool. J Mol Biol 215, 403–410 (1990).

53. Specificity of the Hairpin Ribozyme: SEQUENCE REQUIREMENTS SURROUNDING THE CLEAVAGE SITE. Journal of Biological Chemistry 274, 29376–29380 (1999).

54. Website. https://doi.org/10.1101/2023.09.30.560155 doi:10.1101/2023.09.30.560155.

55. Vaish, N. K., Kore, A. R. & Eckstein, F. Recent developments in the hammerhead ribozyme field. Nucleic Acids Res 26, 5237–5242 (1998).

56. Fornace, M. E. et al. NUPACK: Analysis and Design of Nucleic Acid Structures, Devices, and Systems. (2022) doi:10.26434/chemrxiv-2022-xv98l.

57. Website. https://doi.org/10.1093/molbev/msu300?force_isolation=true doi:10.1093/molbev/msu300?force_isolation=true.

58. Website. https://doi.org/10.1093/molbev/msw046 doi:10.1093/molbev/msw046.

59. FARFAR2: Improved De Novo Rosetta Prediction of Complex Global RNA Folds. Structure 28, 963–976.e6 (2020).

60. Pettersen, E. F. et al. UCSF Chimera--a visualization system for exploratory research and analysis. J Comput Chem 25, 1605–1612 (2004).

