## Supplementary figure 3 for "A food chain of ribozymes built on an ancestral strand"

| | C | C | C | C | C | HHR | HHR | HHR |
|---|---|---|---|---|---|---|---|---|
| Eng | 1 | | | | | 1 | 1 | 1 |
| Sub H | | | 25 | 55 | 56 | 25 | 55 | 56 |
| Leg H | | 47 | | | | 47 | 47 | 47 |

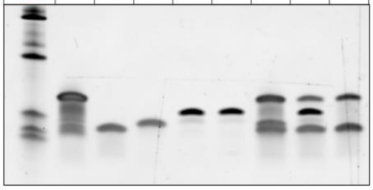


| | C | C | C | C | C | HHR | HHR | HHR |
|---|---|---|---|---|---|---|---|---|
| Eng | 1 | | | | | 1 | 1 | 1 |
| Sub H | | | 25 | 55 | 56 | 25 | 55 | 56 |
| Leg H | | 54 | | | | 54 | 54 | 54 |

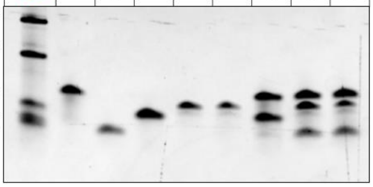


| | C | C | C | C | C | HHR | HHR | HHR |
|---|---|---|---|---|---|---|---|---|
| Eng | 1 | | | | | 1 | 1 | 1 |
| Sub H | | | 57 | 58 | 59 | 57 | 58 | 59 |
| Leg H | | 54 | | | | 54 | 54 | 54 |

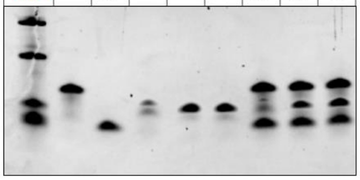


| | C | C | C | C | C | HHR | HHR | HHR |
|---|---|---|---|---|---|---|---|---|
| Eng | 1 | | | | | 1 | 1 | 1 |
| Sub H | | | 57 | 58 | 59 | 57 | 58 | 59 |
| Leg H | | 47 | | | | 47 | 47 | 47 |

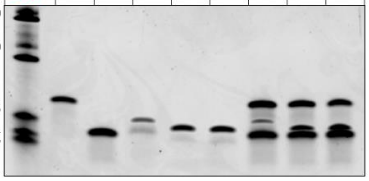


| | C | C | C | C | C | HHR | HHR | HHR |
|---|---|---|---|---|---|---|---|---|
| Eng | 1 | | | | | 1 | 1 | 1 |
| Sub H | | | 25 | 55 | 56 | 25 | 55 | 56 |
| Leg H | | 51 | | | | 51 | 51 | 51 |

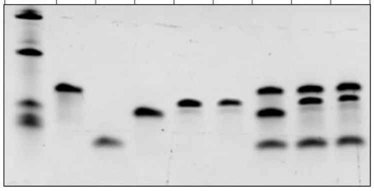


| | C | C | C | C | C | HHR | HHR | HHR |
|---|---|---|---|---|---|---|---|---|
| Eng | 1 | | | | | 1 | 1 | 1 |
| Sub H | | | 57 | 58 | 59 | 57 | 58 | 59 |
| Leg H | | 51 | | | | 51 | 51 | 51 |

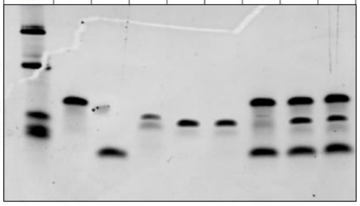


| | C | C | C | C | C | HHR | HHR | HHR |
|---|---|---|---|---|---|---|---|---|
| Eng | 1 | | | | | 1 | 1 | 1 |
| Sub H | | | 25 | 55 | 56 | 25 | 55 | 56 |
| Leg H | | 39 | | | | 39 | 39 | 39 |

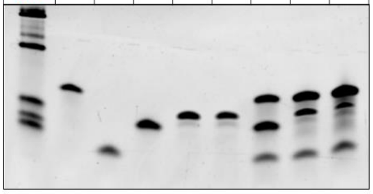


| | C | C | C | C | C | HHR | HHR | HHR |
|---|---|---|---|---|---|---|---|---|
| Eng | 1 | | | | | 1 | 1 | 1 |
| Sub H | | | 25 | 55 | 56 | 25 | 55 | 56 |
| Leg H | | 53 | | | | 53 | 53 | 53 |

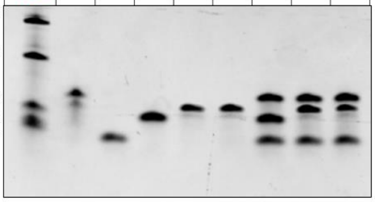


| | C | C | C | C | C | HHR | HHR | HHR |
|---|---|---|---|---|---|---|---|---|
| Eng | 1 | | | | | 1 | 1 | 1 |
| Sub H | | | 57 | 58 | 59 | 57 | 58 | 59 |
| Leg H | | 53 | | | | 53 | 53 | 53 |

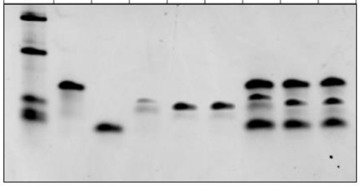


| | C | C | C | C | C | HHR | HHR | HHR |
|---|---|---|---|---|---|---|---|---|
| Eng | 1 | | | | | 1 | 1 | 1 |
| Sub H | | | 57 | 58 | 59 | 57 | 58 | 59 |
| Leg H | | 39 | | | | 39 | 39 | 39 |

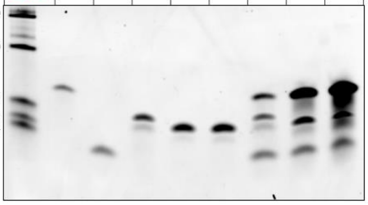


| | C | C | C | C | C | HHR | HHR | HHR |
|---|---|---|---|---|---|---|---|---|
| Eng | 1 | | | | | 1 | 1 | 1 |
| Sub H | | | 25 | 55 | 56 | 25 | 55 | 56 |
| Leg H | | 10 | | | | 10 | 10 | 10 |

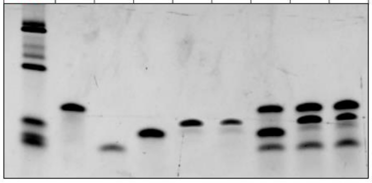


| | C | C | C | C | C | HHR | HHR | HHR |
|---|---|---|---|---|---|---|---|---|
| Eng | 1 | | | | | 1 | 1 | 1 |
| Sub H | | | 57 | 58 | 59 | 57 | 58 | 59 |
| Leg H | | 10 | | | | 10 | 10 | 10 |

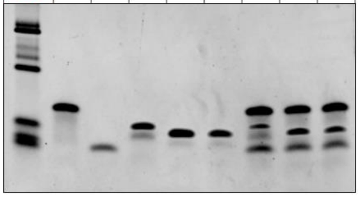


| | C | C | C | C | C | HHR | HHR | HHR |
|---|---|---|---|---|---|---|---|---|
| Eng | 1 | | | | | 1 | 1 | 1 |
| Sub H | | | 25 | 55 | 56 | 25 | 55 | 56 |
| Leg H | | 38 | | | | 38 | 38 | 38 |

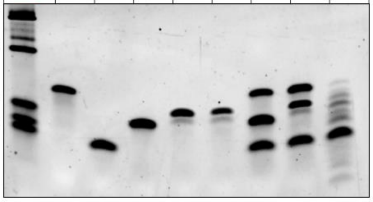


| | C | C | C | C | C | HHR | HHR | HHR |
|---|---|---|---|---|---|---|---|---|
| Eng | 1 | | | | | 1 | 1 | 1 |
| Sub H | | | 25 | 55 | 56 | 25 | 55 | 56 |
| Leg H | | 50 | | | | 50 | 50 | 50 |

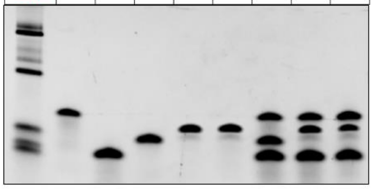


| C | C | C | C | C | HHR | HHR | HHR | |
|---|---|---|---|---|---|---|---|---|
| 1 | | | | | 1 | 1 | 1 | Eng |
| | | 57 | 58 | 59 | 57 | 58 | 59 | Sub H |
| | 50 | | | | 50 | 50 | 50 | Leg H |

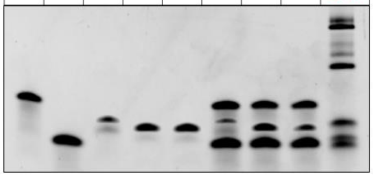


| | C | C | C | C | C | HHR | HHR | HHR |
|---|---|---|---|---|---|---|---|---|
| Eng | 1 | | | | | 1 | 1 | 1 |
| Sub H | | | 57 | 58 | 59 | 57 | 58 | 59 |
| Leg H | | 38 | | | | 38 | 38 | 38 |

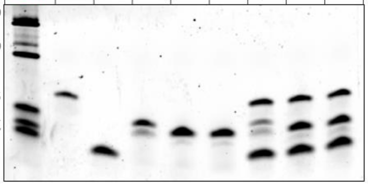


| | C | C | C | C | C | HHR | HHR | HHR |
|---|---|---|---|---|---|---|---|---|
| Eng | 1 | | | | | 1 | 1 | 1 |
| Sub H | | | 25 | 55 | 56 | 25 | 55 | 56 |
| Leg H | | 49 | | | | 49 | 49 | 49 |

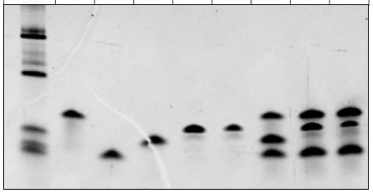


| | C | C | C | C | C | HHR | HHR | HHR |
|---|---|---|---|---|---|---|---|---|
| Eng | 1 | | | | | 1 | 1 | 1 |
| Sub H | | | 57 | 58 | 59 | 57 | 58 | 59 |
| Leg H | | 49 | | | | 49 | 49 | 49 |

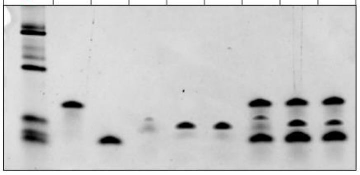

| | C | C | C | C | C | HHR | HHR | HHR |
|---|---|---|---|---|---|---|---|---|
| Eng | 1 | | | | | 1 | 1 | 1 |
| Sub H | | | 25 | 55 | 56 | 25 | 55 | 56 |
| Leg H | | 43 | | | | 43 | 43 | 43 |

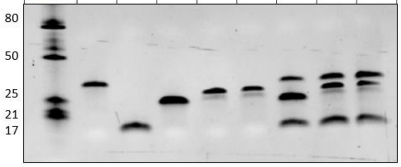


| | C | C | C | C | C | HHR | HHR | HHR |
|---|---|---|---|---|---|---|---|---|
| Eng | 1 | | | | | 1 | 1 | 1 |
| Sub H | | | 25 | 55 | 56 | 25 | 55 | 56 |
| Leg H | | 18 | | | | 18 | 18 | 18 |

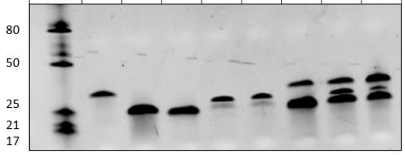


| | C | C | C | C | C | HHR | HHR | HHR |
|---|---|---|---|---|---|---|---|---|
| Eng | 1 | | | | | 1 | 1 | 1 |
| Sub H | | | 57 | 58 | 59 | 57 | 58 | 59 |
| Leg H | | 18 | | | | 18 | 18 | 18 |

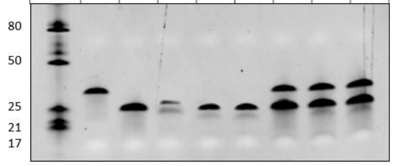


| | C | C | C | C | C | HHR | HHR | HHR |
|---|---|---|---|---|---|---|---|---|
| Eng | 1 | | | | | 1 | 1 | 1 |
| Sub H | | | 57 | 58 | 59 | 57 | 58 | 59 |
| Leg H | | 43 | | | | 43 | 43 | 43 |

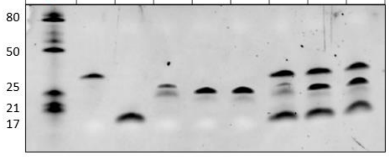


| | C | C | C | C | C | HHR | HHR | HHR |
|---|---|---|---|---|---|---|---|---|
| Eng | 1 | | | | | 1 | 1 | 1 |
| Sub H | | | 25 | 55 | 56 | 25 | 55 | 56 |
| Leg H | | 48 | | | | 48 | 48 | 48 |

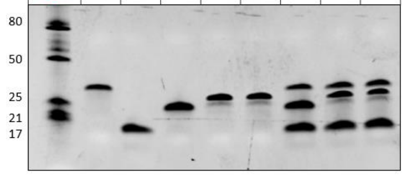


| | C | C | C | C | C | HHR | HHR | HHR |
|---|---|---|---|---|---|---|---|---|
| Eng | 1 | | | | | 1 | 1 | 1 |
| Sub H | | | 57 | 58 | 59 | 57 | 58 | 59 |
| Leg H | | 48 | | | | 48 | 48 | 48 |

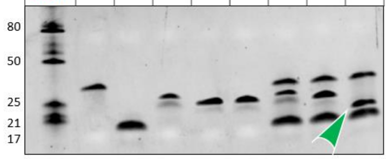


| | C | C | C | C | C | HHR | HHR | HHR |
|---|---|---|---|---|---|---|---|---|
| Eng | 1 | | | | | 1 | 1 | 1 |
| Sub H | | | 25 | 55 | 56 | 25 | 55 | 56 |
| Leg H | | 11 | | | | 11 | 11 | 11 |

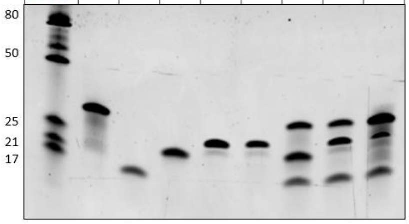


| | C | C | C | C | C | HHR | HHR | HHR |
|---|---|---|---|---|---|---|---|---|
| Eng | 1 | | | | | 1 | 1 | 1 |
| Sub H | | | 25 | 55 | 56 | 25 | 55 | 56 |
| Leg H | | 46 | | | | 46 | 46 | 46 |

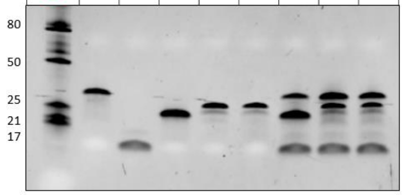


| | C | C | C | C | C | HHR | HHR | HHR |
|---|---|---|---|---|---|---|---|---|
| Eng | 1 | | | | | 1 | 1 | 1 |
| Sub H | | | 57 | 58 | 59 | 57 | 58 | 59 |
| Leg H | | 46 | | | | 46 | 46 | 46 |

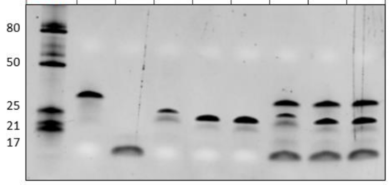


| | C | C | C | C | C | HHR | HHR | HHR |
|---|---|---|---|---|---|---|---|---|
| Eng | 1 | | | | | 1 | 1 | 1 |
| Sub H | | | 57 | 58 | 59 | 57 | 58 | 59 |
| Leg H | | 11 | | | | 11 | 11 | 11 |

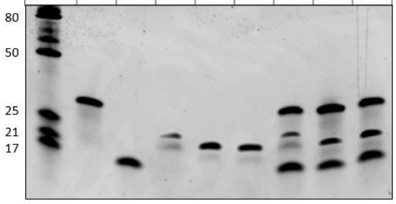


| | C | C | C | C | C | HHR | HHR | HHR |
|---|---|---|---|---|---|---|---|---|
| Eng | 1 | | | | | 1 | 1 | 1 |
| Sub H | | | 25 | 55 | 56 | 25 | 55 | 56 |
| Leg H | | 21 | | | | 21 | 21 | 21 |

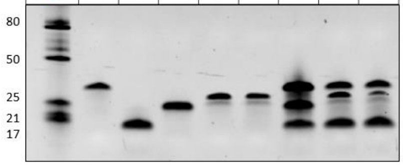


| | C | C | C | C | C | HHR | HHR | HHR |
|---|---|---|---|---|---|---|---|---|
| Eng | 1 | | | | | 1 | 1 | 1 |
| Sub H | | | 57 | 58 | 59 | 57 | 58 | 59 |
| Leg H | | 21 | | | | 21 | 21 | 21 |

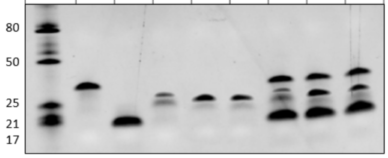


| | C | C | C | C | C | HHR | HHR | HHR |
|---|---|---|---|---|---|---|---|---|
| Eng | 1 | | | | | 1 | 1 | 1 |
| Sub H | | | 57 | 58 | 59 | 57 | 58 | 59 |
| Leg H | | 52 | | | | 52 | 52 | 52 |

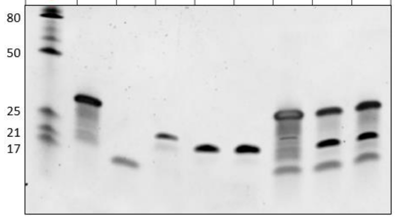


| | C | C | C | C | C | HHR | HHR | HHR |
|---|---|---|---|---|---|---|---|---|
| Eng | 1 | | | | | 1 | 1 | 1 |
| Sub H | | | 25 | 55 | 56 | 25 | 55 | 56 |
| Leg H | | 45 | | | | 45 | 45 | 45 |

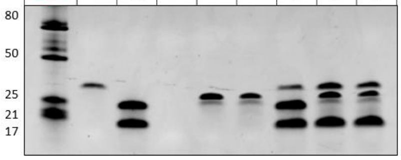


| | C | C | C | C | C | HHR | HHR | HHR |
|---|---|---|---|---|---|---|---|---|
| Eng | 1 | | | | | 1 | 1 | 1 |
| Sub H | | | 57 | 58 | 59 | 57 | 58 | 59 |
| Leg H | | 45 | | | | 45 | 45 | 45 |

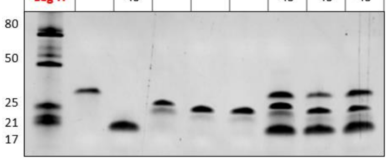


| | C | C | C | C | C | HHR | HHR | HHR |
|---|---|---|---|---|---|---|---|---|
| Eng | 1 | | | | | 1 | 1 | 1 |
| Sub H | | | 25 | 55 | 56 | 25 | 55 | 56 |
| Leg H | | 52 | | | | 52 | 52 | 52 |

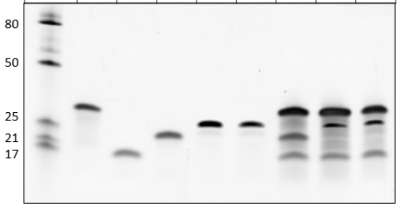


| | C | C | C | C | C | HHR | HHR | HHR |
|---|---|---|---|---|---|---|---|---|
| Eng | 1 | | | | | 1 | 1 | 1 |
| Sub H | | | 25 | 55 | 56 | 25 | 55 | 56 |
| Leg H | | 44 | | | | 44 | 44 | 44 |

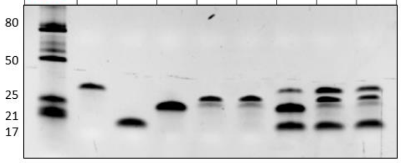


| | C | C | C | C | C | HHR | HHR | HHR |
|---|---|---|---|---|---|---|---|---|
| Eng | 1 | | | | | 1 | 1 | 1 |
| Sub H | | | 57 | 58 | 59 | 57 | 58 | 59 |
| Leg H | | 44 | | | | 44 | 44 | 44 |

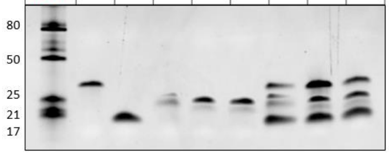

| | C | C | C | C | C | C | 2C | 2C | 2C | HPR | HHR |
|---|---|---|---|---|---|---|---|---|---|---|---|
| Eng | 1 | | | | | 4 | 1 | 1 | | 1 | 4 |
| Leg P / Sub H | | 14 | | | 15 | | 14 | | 14 | 14 | 15 |
| Leg H | | | | 10 | | | | | | | 10 |
| Sub P | | | 18 | | | | | 18 | 18 | 18 | |

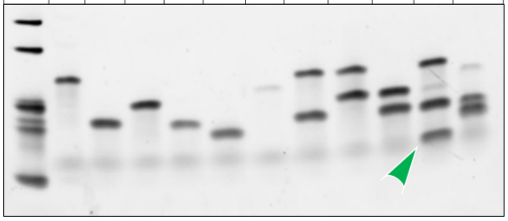


| | C | C | C | C | C | C | C | HPR | HHR | HPR | HHR |
|---|---|---|---|---|---|---|---|---|---|---|---|
| Eng | 1 | | | | | | 20 | 1 | 20 | 1 | 20 |
| Leg P / Sub H | | | 28 | 35 | | | | 28 | 28 | 35 | 28 |
| Leg H | | | | | 22 | 23 | | | 22 | | 23 |
| Sub P | | 19 | | | | | | 19 | | 19 | |

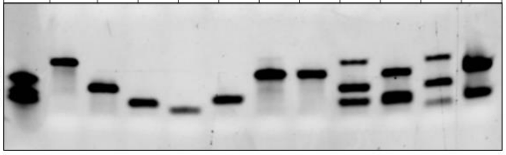


| | C | C | C | C | C | C | HPR | HHR | HPR |
|---|---|---|---|---|---|---|---|---|---|
| Eng | 20 | | | | | | 20 | 20 | 20 |
| Leg P / Sub H | | | 30 | 33 | 31 | | 31 | 30 | 33 |
| Leg H | | 23 | | | | | | 23 | |
| Sub P | | | | | | 36 | 36 | | 36 |

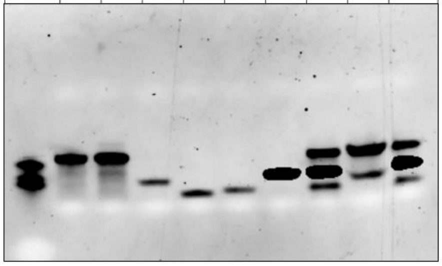


| | C | C | C | C | 2C | 2C | 2C | HHR | HHR |
|---|---|---|---|---|---|---|---|---|---|
| Eng | | | 4 | | | 4 | 4 | 4 | 4 |
| Sub H | 16 | | | | 16 | | 16 | 16 | 16 |
| Leg H | | 11 | | 8 | 11 | 11 | | 11 | 8 |

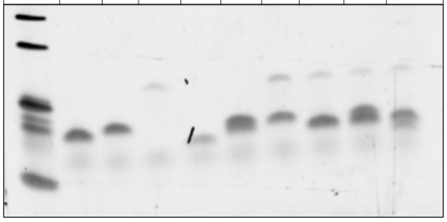


| | C | C | C | C | C | HPR | HPR | HPR |
|---|---|---|---|---|---|---|---|---|
| Eng | 20 | | | | | 20 | 20 | 20 |
| Leg P | | 34 | | 28 | 27 | 27 | 34 | 28 |
| Sub P | | | 36 | | | 36 | 36 | 36 |

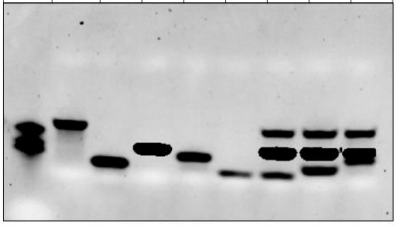


| | C | C | C | C | C | HHR | HHR | HHR |
|---|---|---|---|---|---|---|---|---|
| Eng | 1 | | | | | 1 | 1 | 1 |
| Sub H | | | 25 | 55 | 56 | 25 | 55 | 56 |
| Leg H | | 42 | | | | 42 | 42 | 42 |

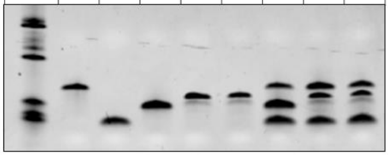


| | C | C | C | C | C | HHR | HHR | HHR |
|---|---|---|---|---|---|---|---|---|
| Eng | 1 | | | | | 1 | 1 | 1 |
| Sub H | | | 25 | 55 | 56 | 25 | 55 | 56 |
| Leg H | | 41 | | | | 41 | 41 | 41 |

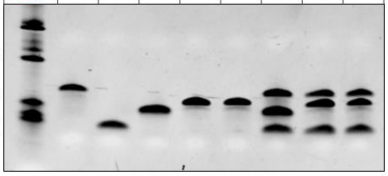


| | C | C | C | C | C | HHR | HHR | HHR |
|---|---|---|---|---|---|---|---|---|
| Eng | 1 | | | | | 1 | 1 | 1 |
| Sub H | | | 25 | 55 | 56 | 25 | 55 | 56 |
| Leg H | | 40 | | | | 40 | 40 | 40 |

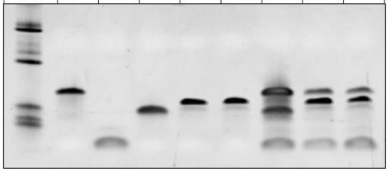


| | C | C | C | C | C | HHR | HHR | HHR |
|---|---|---|---|---|---|---|---|---|
| Eng | 1 | | | | | 1 | 1 | 1 |
| Sub H | | | 25 | 55 | 56 | 25 | 55 | 56 |
| Leg H | | 5 | | | | 5 | 5 | 5 |

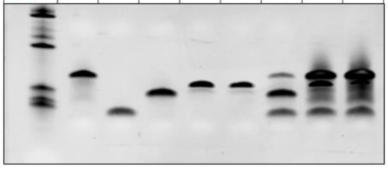


| | C | C | C | C | C | 2C | HHR | HHR |
|---|---|---|---|---|---|---|---|---|
| Eng | 1 | | | | | 1 | 1 | 1 |
| Sub H | | | 25 | 55 | 56 | | 55 | 56 |
| Leg H | | 37 | | | | 37 | 37 | 37 |

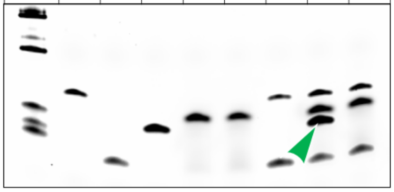


| | C | C | C | C | C | HPR | HPR | HPR |
|---|---|---|---|---|---|---|---|---|
| Eng | 1 | | | | | 1 | 1 | 1 |
| Leg P | | | 26 | 29 | 32 | 32 | 26 | 29 |
| Sub P | | 19 | | | | 19 | 19 | 19 |

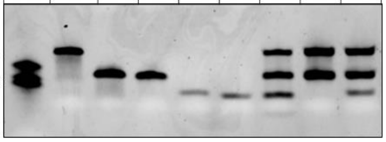


| | C | C | C | C | C | HHR | HHR | HHR |
|---|---|---|---|---|---|---|---|---|
| Eng | 1 | | | | | 1 | 1 | 1 |
| Sub H | | | 57 | 58 | 59 | 57 | 58 | 59 |
| Leg H | | 42 | | | | 42 | 42 | 42 |

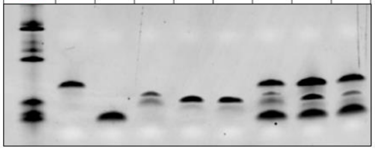


| | C | C | C | C | C | HHR | HHR | HHR |
|---|---|---|---|---|---|---|---|---|
| Eng | 1 | | | | | 1 | 1 | 1 |
| Sub H | | | 57 | 58 | 59 | 57 | 58 | 59 |
| Leg H | | 41 | | | | 41 | 41 | 41 |

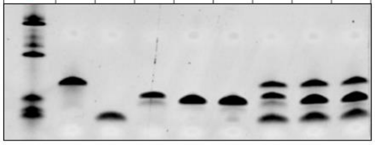


| | C | C | C | C | C | HHR | HHR | HHR |
|---|---|---|---|---|---|---|---|---|
| Eng | 1 | | | | | 1 | 1 | 1 |
| Sub H | | | 57 | 58 | 59 | 57 | 58 | 59 |
| Leg H | | 40 | | | | 40 | 40 | 40 |

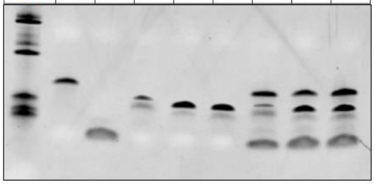


| | C | C | C | C | C | HHR | HHR | HHR |
|---|---|---|---|---|---|---|---|---|
| Eng | 1 | | | | | 1 | 1 | 1 |
| Sub H | | | 57 | 58 | 59 | 57 | 58 | 59 |
| Leg H | | 5 | | | | 5 | 5 | 5 |

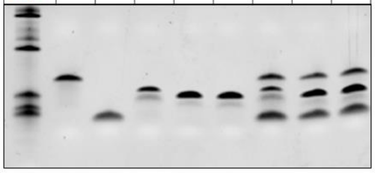


| | C | C | C | C | C | HHR | HHR | HHR |
|---|---|---|---|---|---|---|---|---|
| Eng | 1 | | | | | 1 | 1 | 1 |
| Sub H | | | 57 | 58 | 59 | 57 | 58 | 59 |
| Leg H | | 37 | | | | 37 | 37 | 37 |

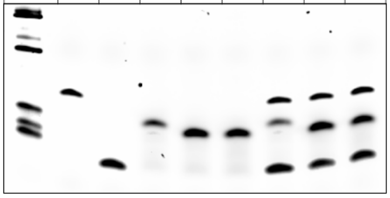


| | C | C | C | HHR | C | C | C | HHR |
|---|---|---|---|---|---|---|---|---|
| Eng | | | 20 | 20 | 1 | | | 1 |
| Sub H | 25 | | | 25 | | | 34 | 34 |
| Leg H | | 21 | | 21 | | 24 | | 24 |

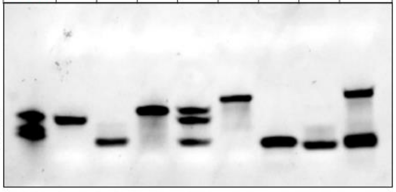

| | C | C | C | C | C | C | HHR | HPR | HHR | HHR |
|---|---|---|---|---|---|---|---|---|---|---|
| Eng | | | 1 | | | | 1 | 1 | 1 | 1 |
| Leg P / Sub H | 14 | | | | | 16 | 14 | 14 | 14 | 16 |
| Leg H | | 5 | | | 12 | | 5 | | 12 | 5 |
| Sub P | | | | 18 | | | | 18 | | |

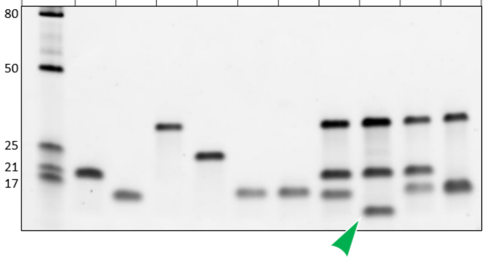


| | | | | | | | | HPR | HPR | HPR | HPR |
|---|---|---|---|---|---|---|---|---|---|---|---|
| Eng | 3 | | | | 1 | 2 | | 2 | 3 | 1 | 2 |
| Leg P | | 13 | | | | | 14 | 13 | 13 | 13 | 14 |
| Sub P | | | 19 | 18 | | | | 18 | 19 | 18 | 18 |

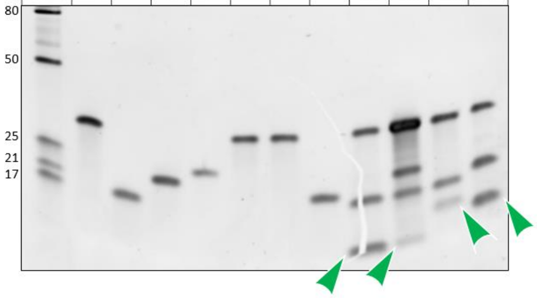


| | C | C | C | C | C | C | HHR | HHR | HHR | HHR |
|---|---|---|---|---|---|---|---|---|---|---|
| Eng | | | 4 | | | | 4 | 4 | 4 | 4 |
| Sub H | 16 | | | | | | 16 | 16 | 16 | 16 |
| Leg H | | 6 | | 9 | 5 | 7 | 6 | 9 | 5 | 7 |

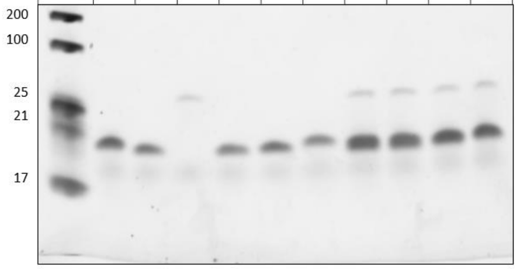


| | C | C | C | C | C | C | HHR | HHR | HHR | HHR |
|---|---|---|---|---|---|---|---|---|---|---|
| Eng | | | | 2 | 1 | | 1 | 2 | 2 | 2 |
| Sub H | 16 | | | | | | 16 | 16 | 16 | 16 |
| Leg H | | 6 | 7 | | | 8 | 6 | 6 | 7 | 8 |

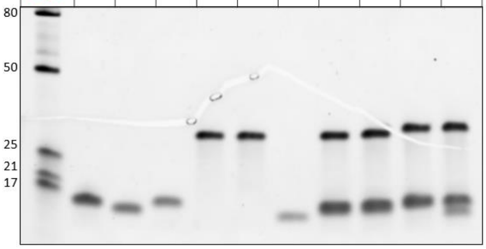


| | C | C | C | C | C | C | HHR | HHR | HHR |
|---|---|---|---|---|---|---|---|---|---|
| Eng | | | 1 | | 2 | 4 | 1 | 2 | 4 |
| Sub H | 15 | | | | | | 15 | 15 | 15 |
| Leg H | | 12 | | 10 | | | 12 | 10 | 10 |

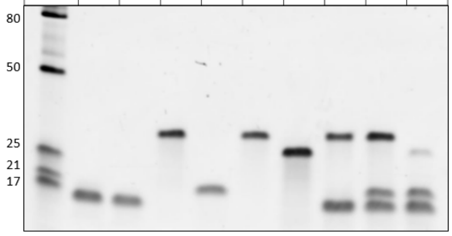


| | C | C | C | C | C | C | HHR | HHR |
|---|---|---|---|---|---|---|---|---|
| Eng | | | 2 | | 4 | | 2 | 2 |
| Sub H | 16 | | | | | | 16 | 16 |
| Leg H | | 9 | | 11 | | 8 | 9 | 11 |

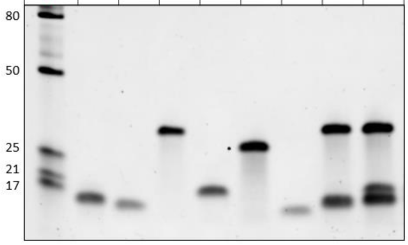
