## Supplementary figures and images for "A food chain of ribozymes built on an ancestral strand"

### Supplementary figure 1

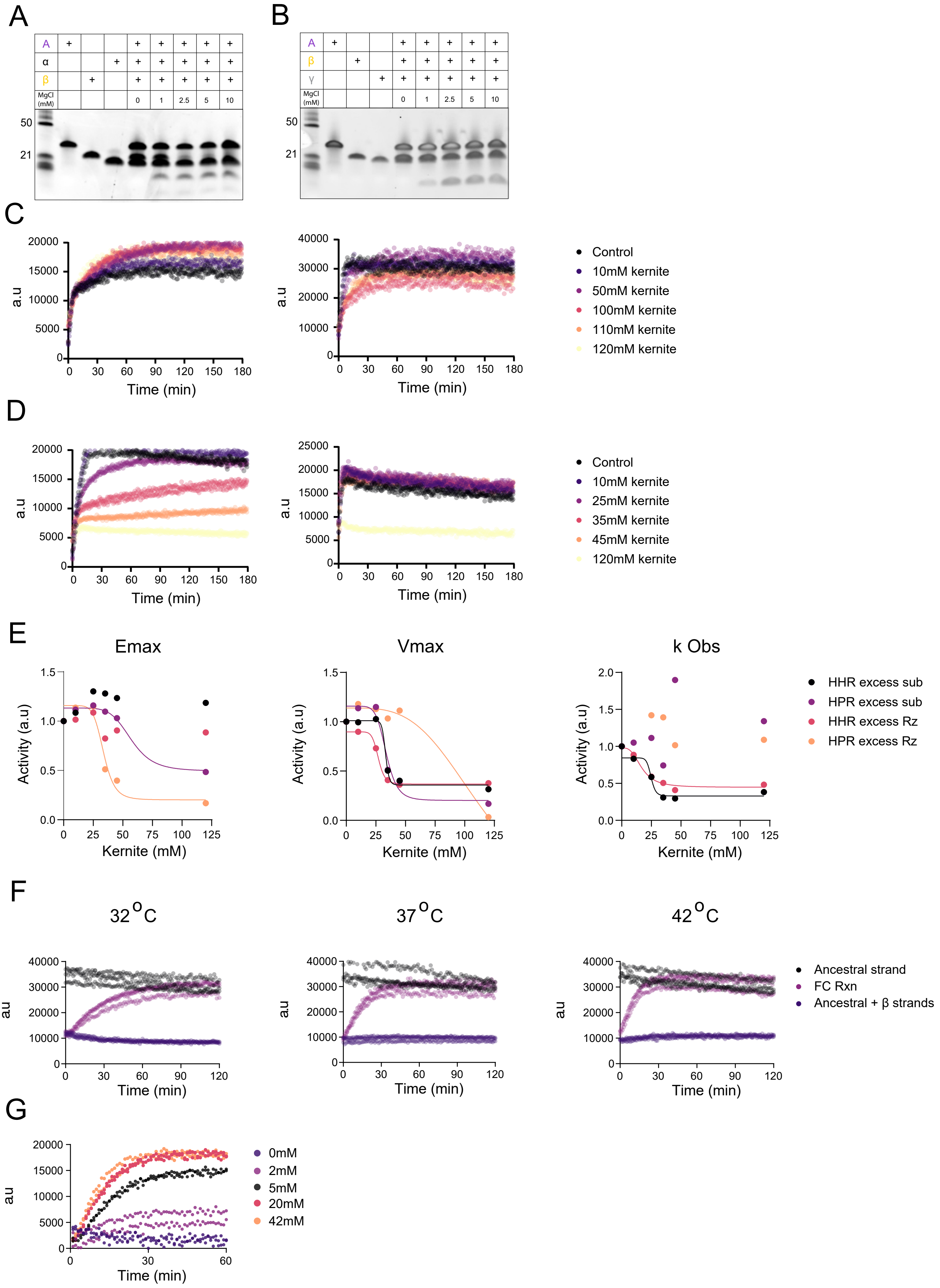

### Supplementary figure 2

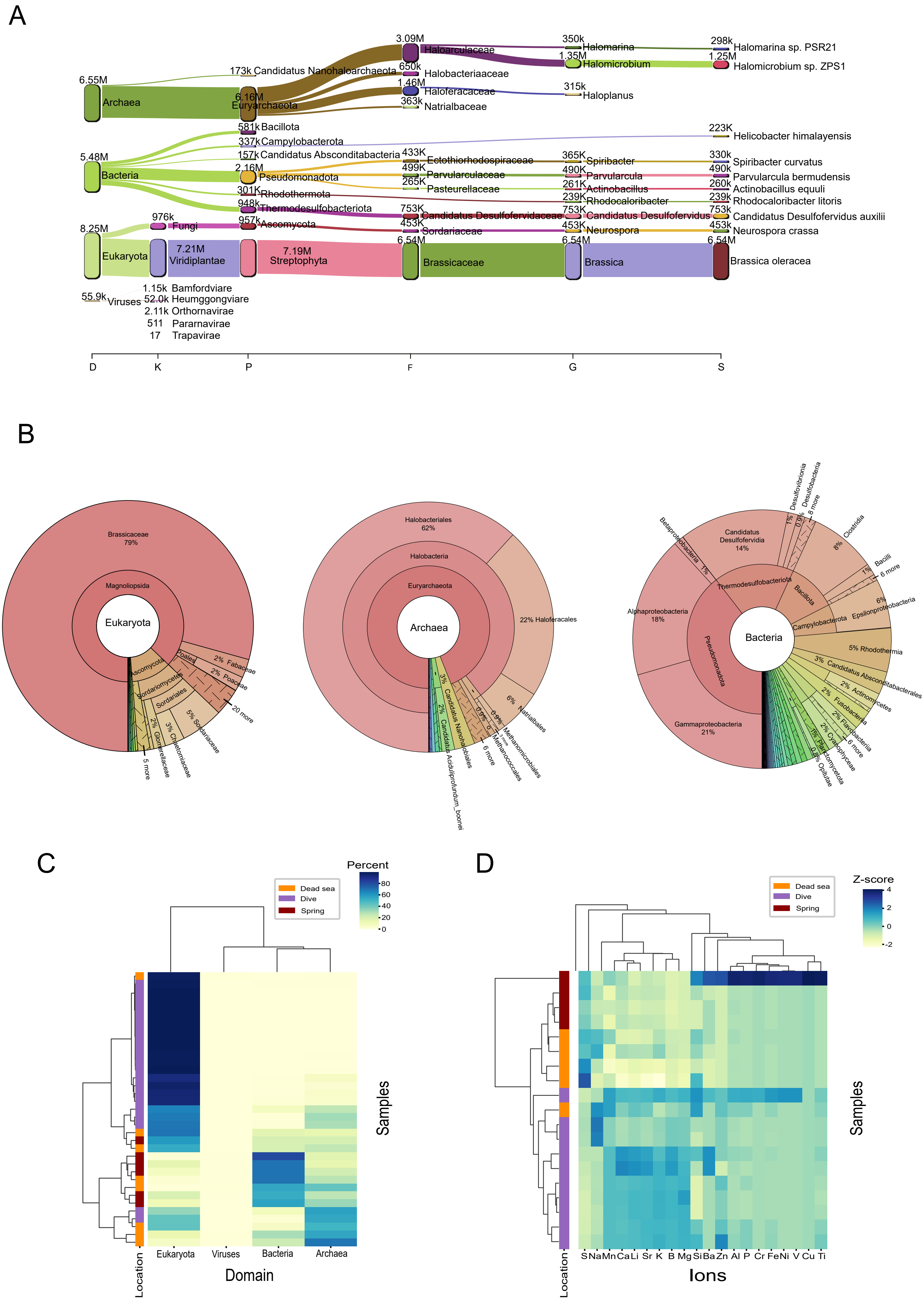
